# Functional genomics reveals mediators of beta cell survival in ER stress and type 2 diabetes risk

**DOI:** 10.64898/2026.03.30.715154

**Authors:** Mei-Lin Okino, Han Zhu, Sierra Corban, Paola Benaglio, Leyan Huang, Jovina Djulamsah, Rebecca Tseng, Bailey O’Mahony, Kennedy Vanderstel, Ruth Elgamal, Michael Miller, Allen Wang, Maike Sander, Kyle Gaulton

**Author notes:** Authors equally contributed to this work (Mei-Lin Okino, Han Zhu). Authors jointly supervised this work (Han Zhu, Maike Sander, Kyle Gaulton). Correspondence to: Kyle Gaulton.

## Abstract

Endoplasmic reticulum (ER) stress in pancreatic beta cells contributes to impaired function and type 2 diabetes (T2D). In this study we used genome-wide perturbation screens and genomic profiling in beta cells to identify novel mediators of ER stress responses and diabetes risk. We defined gene regulatory networks in beta cells and identified specific networks both enriched for T2D risk variants and with altered expression in ER stress. We performed a genome-wide loss-of-function CRISPR screen for survival under ER stress in EndoC-bH1 cells, which identified 214 beta cell survival genes many of which were novel and mapped in pathways related to insulin secretion, mitochondrial and ER-Golgi transport, and post-translational modification. Beta cell survival genes collectively had limited genomic change in stress yet showed significant, independent enrichment for T2D risk variants, including novel T2D candidate gene *DTNB* which we validated promotes beta cell survival during stress. Overall, our results revealed mediators of ER stress responses in beta cells and identified new therapeutic targets to preserve beta cells in diabetes pathogenesis.

## Introduction

Pancreatic beta cells play a critical role in glucose homeostasis and metabolism, and islet dysfunction is a major contributor to the development of type 2 diabetes (T2D)^1,2^. Due to the high demands of insulin production, beta cells are susceptible to the protein-folding capacity of the endoplasmic reticulum (ER) being overwhelmed which can lead to ER stress and activation of the unfolded protein response (UPR)^3^. The UPR response is initiated by IRE1a, PERK, and ATF6, which initiates a signaling cascade aimed at restoring ER homeostasis by halting translation and upregulating protein-folding chaperones^4^. Prolonged activation of the UPR through unresolved ER stress impairs insulin production and activates pro-apoptotic pathways^5^. In the context of T2D progression, increased demands for insulin production and secretion coupled with glucolipotoxicity due to elevated glucose and lipid levels can drive chronic ER stress and UPR activation leading to beta cell dysfunction and death^6,7^.

Given the critical importance of ER stress responses in beta cells to T2D, identifying mediators of beta cell ER stress responses may reveal mechanistic insight into how to protect beta cells in T2D progression. Thapsigargin (Tgn) is a small molecule that induces ER stress through inhibition of the sarco/endoplasmic reticulum Ca²⁺-ATPase (SERCA) pump and has been widely utilized as an *in vitro* model of ER stress in beta cells^8,9,10,11^. Recent studies have performed bulk and single cell genomics profiling of pancreatic islets in Tgn-treated cells, which revealed key genes and transcriptional regulators with altered expression under ER stress^12,13^. These studies have also uncovered genes altered under stress that may represent candidate genes at genetic risk loci for T2D. Genomic changes under stress, however, are not necessarily correlated with direct effects on cellular function nor describe which specific functions are affected. Furthermore, genes that play a critical role in response to stress may not always involve direct epigenomic or transcriptomic changes and may instead, for example, consist of changes in protein translation or stability or alternately not require any changes to play a functional role in stress responses.

High-throughput functional screens for example using CRISPR/Cas9 can be utilized to systematically link genes and genomic regions to downstream phenotypes. In beta cells, genome-wide knockout screens using CRISPR have been employed to identify genes modulating survival under inflammatory cytokine stress and to identify genes affecting insulin content^14,15^, which have revealed novel targets to promote beta cell function and have provided functional context to candidate genes involved in genetic risk of diabetes. Functional screens of beta cells, however, not to date been applied to models of ER stress. In this study, we therefore combined genome-wide functional screens with single cell profiling and T2D association data to understand mediators of ER stress and T2D risk in beta cells. Our results reveal genes critical to survival under ER stress, including many not linked to genomic changes under ER stress, and identify new targets to prevent beta cell death during ER stress that may protect against T2D.

## Results

### ER stress-induced genomic changes in pancreatic islets

We determined the genomic response of pancreatic islets to ER stress as well as, for comparison, to inflammatory cytokine stress. We exposed islets from three non-diabetic donors to thapsigargin (Tgn), pro-inflammatory cytokines (IL-1Β, IFN-γ, TNF-α), or no treatment (**Table S1**), and then performed bulk ATAC-seq and RNA-seq. Overall, Tgn and cytokines induced widespread, distinct changes to the islet epigenome and transcriptome compared to untreated cells **(Figure 1B).** Differential gene expression analysis revealed thousands of genes with significant changes (FDR<0.05) in expression in Tgn and cytokine treated islets **(Figure 1C, Table S2)**. ER and cytokine stress up-regulated different sets of genes; for example, cytokines upregulated chemokine and inflammatory signaling genes such as *LRRK2*^16^, *CX3CL1*^17^*, and JAK2*^18^ (log2(fold change[FC])=0.72, 5.54, 1.89; FDR<.05) (**Figure S1A, Table S2**) while Tgn up-regulated genes associated with ER stress such as *ATF3,* a TF involved in the regulation of apoptosis^19^, and *TMBIM6*^20^, a positive autophagy regulator (log2(FC)=2.09, 0.52; FDR<.05) **(Figure S1A, Table S2**).

**Figure 1.**
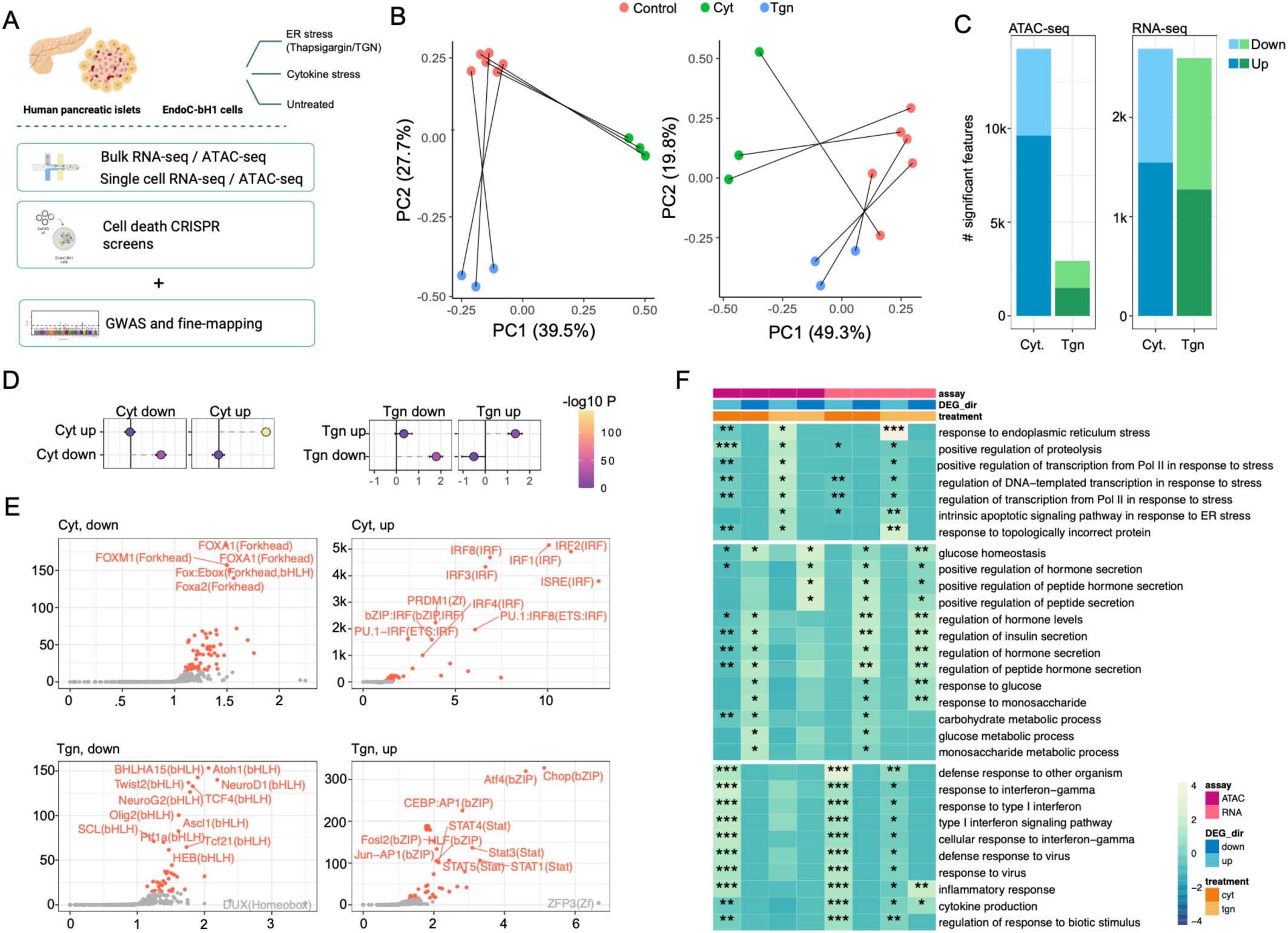
Epigenomic and transcriptomic responses to ER stress and inflammation in pancreatic islets. (A) Study overview schematic. (B) Principal component analysis (PCA) of cCRE accessibility (right) and gene expression (left). TGN- or cytokine-treated samples are connected to controls from the same islet donor. (C) Bar plots of significantly differentially accessible cCREs (left) and differentially expressed genes (right). Differential expression and accessibility are defined at FDR<.05. (D) Enrichment (Fisher’s exact test) for differentially up- or down-regulated genes in the set of closest genes to differentially up- or down-regulated cCREs**. (**E) Volcano plots of known motifs enriched in up- and down-regulated cCREs. Fold change is calculated as the expected divided by observed number of cCREs containing a given motif. P-values are from HOMER. (F) Enrichment for selected pathways from gene ontology in sets of up- and down-regulated genes and cCREs. Enrichment p-values were calculated using Goseq for gene analysis and GREAT for cCRE analysis, respectively. Color values are scaled by sample. **FDR<1E-05; *FDR<0.05

We defined 196,298 candidate cis-regulatory elements (cCREs) using ATAC-seq profiles across samples and conditions and performed differential analyses of cCRE activity in ER and inflammatory stress. There were thousands of cCREs with differential activity (FDR<.05) in stress including 2,938 ER stress-responsive cCREs and 14,258 cytokine-responsive cCREs **(Figure 1C, Table S3)**. Stress-responsive cCREs showed concordant direction of effect with the expression of proximal genes **(Figure 1D, Table S5**). For example, a cCRE up-regulated in ER stress (log2(fold change[FC])=1.14, FDR<.05) (**Table S3, Figure S1B)** mapped to an intron of *ATF3,* which had up-regulated expression (**Table S3),** and cCREs up-regulated in inflammatory stress mapped to the up-regulated genes *LRRK2, CX3CL1,* and *JAK2* **(Figure S1B, Table S2-3**). TF motif analysis revealed that ER-stress responsive cCREs were enriched (FDR<.05) for motifs for mediators of UPR including ATF4 and CHOP^4^ (**Table S4**), while cCREs down-regulated in ER stress were enriched (FDR<.05) for beta cell TFs such as FOXA2 and NEUROD1^21^ (**Table S4**). Cytokine-responsive cCREs were enriched (FDR<.05) for IRF motifs (**Table S4**), while cCREs down-regulated in cytokines were enriched for similar motifs as ER stress (**Figure 1E, Table S4**). We annotated cREs based on islet chromatin state^22^, and cREs with altered accessibility in ER stress were strongly enriched for enhancer states **(Figure S1C, Table S5**), emphasizing that epigenomic responses during stress in beta cells are primarily driven through enhancer accessibility.

Finally, we determined biological processes affected by ER and inflammatory stress in islets. In ER stress, in both the transcriptome and epigenome, pathways related to ER stress response and apoptotic signaling were significantly (FDR<.05) up-regulated (**Figure 1F; Table S6**). Conversely, in cytokine stress, pathways related to interferon signaling, inflammatory response, and viral response were all significantly up-regulated (FDR<.05) (**Figure 1F, Table S6**). While the pathways up-regulated in inflammatory and ER stress were largely distinct, key pathways related to beta cell function including glucose homeostasis and hormone secretion were significantly down-regulated (FDR<.05) under both treatment conditions (**Figure 1F, Table S6**). Illustrative of this, *SLC30A8,* a gene associated with insulin secretion^23^, had downregulated expression and cCRE activity under both inflammatory and ER stress (log2(FC)=-1.18, -0.89 for Cyt and Tgn, respectively; both FDR<.05) (**Table S2**).

Overall, our findings show that ER stress drives extensive transcriptomic and epigenomic changes in pancreatic islets and that these changes are broadly distinct from inflammatory stress responses.

### Single-nucleus multiomics reveals islet cell type-specific responses to ER stress

To understand the effects of ER stress on islet cell types, we next performed single cell multiome (joint snATAC-seq and RNA-seq) profiling of islets from non-diabetic donors treated with Tgn, cytokines, or control (n=5, 4, 4 per condition, respectively) using the same treatment duration and concentration as for bulk assays. We further incorporated snATAC-seq data from islets from four additional donors (n=2, 2, 4 for Tgn, cytokines, and control, respectively) and single cell profiles from 21 non-diabetic, pre-T2D and T2D donors from a previous study^24^ (**Figure 2A, Figure S2A, Table S1)**. Following quality control involving barcode filtering, correcting for ambient RNA^25^, and doublet removal^26^ (**Figure S2B, Table S7**), we identified 12 clusters which we annotated using marker gene expression of pancreatic cell types (**Figure 2B, Figure S2C**). One cluster expressed both beta and alpha cell marker genes, which we annotated as ‘alpha+beta’, and clustered distinct from other alpha and beta cells in snATAC-seq. As the biological significance of cluster was unclear, we excluded it from further analyses (**Figure S2C-D**).

**Figure 2.**
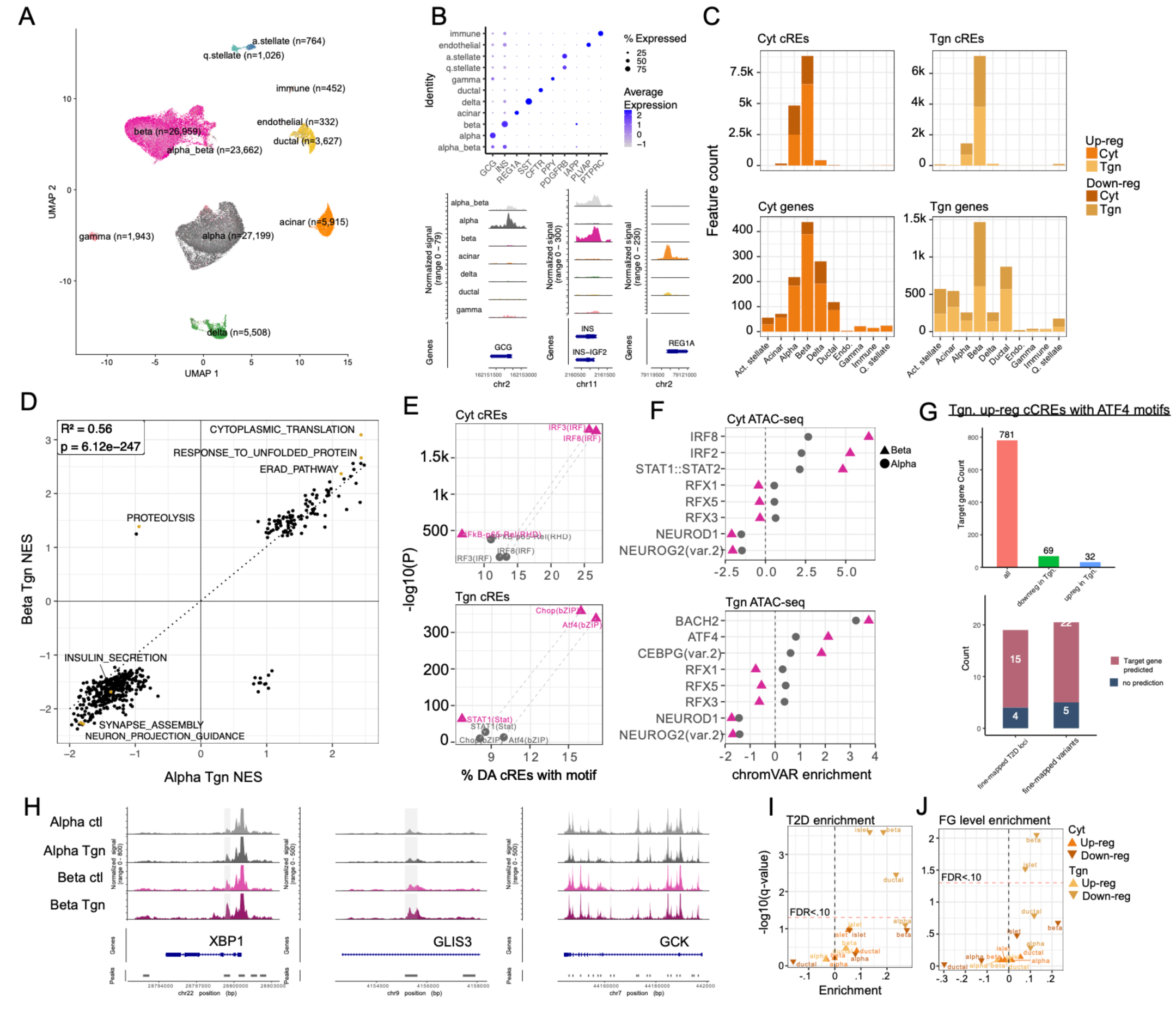
Single nucleus multiomic profiling reveals islet cell type-specific regulatory changes in ER stress. (A) UMAP representation of 128,202 nuclei profiled with single cell multiome (paired snATAC+RNA-seq) or single modality snATAC-seq. Note that “alpha_beta” cells (in light grey) do not form a distinct cluster. (B) Dot plot displaying expression of islet cell type marker genes in islet cell types across treatment conditions (top); genome browser tracks displaying cell type-specific chromatin accessibility at the promoters of cell type marker genes (bottom). (C) Bar plots of significantly differentially accessible cCREs (top) and differentially expressed genes (bottom) in pseudobulk cell types. Differential expression and accessibility are defined at FDR<.05. (D) Scatter plot displaying normalized pathway enrichment scores for alpha and beta cells treated with thapsigargin. (E) Comparison of enrichment for selected motifs in pseudobulk alpha and beta cell cCREs upregulated by exposure to cytokines (top) or TGN (bottom). (F) Comparison of Z-scores representing changes in global accessibility of selected motifs in cytokine-treated (top) or TGN-treated (bottom) pseudobulk alpha and beta cells. (G) Bar plot displaying the total number of predicted target genes of Tgn-upregulated beta cell cCREs with ATF4 motifs, and number of significantly up- and down-regulated genes in beta cells (left); Bar plot displaying the total number of fine-mapped T2D risk loci and variants (from DIAMANTE 2018) in the ATF4 GRN. (F) Genome browser tracks highlighting cCREs with ATF4 motifs at intronic regions of notable genes with pseudobulk beta cell-specific increases in accessibility under ER stress. (G) Enrichment of T2D-associated genes in bulk islet and pseudobulk DEGs as calculated by MAGMA. Enrichment scores are from MAGMA and based on summary statistics from Mahajan et al., *Nature Genetics* (2018). (H) Enrichment of fasting glycemia-associated genes in bulk islet and pseudobulk DEGs as calculated by MAGMA. Summary statistics are from the DIAGRAM consortium^78^.

To evaluate the cell type-specific impact of ER stress and inflammatory stress on gene expression, we performed differential expression analysis in each cell type using DESeq2 (**Figure 2C, Table S8**). Hundreds of genes had significantly altered expression (FDR<.05) in Tgn or cytokine treatment across all cell types. Differentially expressed genes in alpha and beta cells were strongly concordant with changes in bulk islets for both treatments (**Table S9**). We found concordant up-regulation of UPR signaling under ER stress and immune responses under cytokine stress in alpha and beta cells, while pathways related to insulin secretion were downregulated in beta cells in both treatments (**Figure 2D, Table S10**). Interestingly, there was stronger up-regulation of protein degradation and quality control pathways in beta cells compared to alpha cells in stress, including protein catabolism in ER stress (**Figure 2D, Table S10,** Tgn beta NES=1.386, FDR=4.25×10^-4^; Tgn alpha NES=-0.942, FDR=0.83) and ubiquitination in inflammatory stress (**Figure 2D, Table S10:** Cyt beta NES=1.88, FDR=1.18×10^-4^; Cyt alpha NES=1.37, FDR=.20). We also observed pathways associated with neuronal dendrite phenotypes downregulated in ER stress (**Figure 2D, Table S10**), which may indicate impaired insulin secretion dynamics in ER stress given similarities between neurons and beta cells in electrochemical signaling.

We next defined 364,039 total cCREs using snATAC-seq profiles across all cell types. In alpha and beta cells, thousands of cCREs had significant changes (FDR<.05) in accessibility in stress, with few changes in other cell types (**Figure 2C, Table S11**). Stress-responsive cCREs in alpha and beta cells were largely concordant with bulk islets, as well as between themselves (**Table S11**). We performed motif enrichment of stress-responsive cCREs using HOMER^27^. Motifs for IRF TFs were more prominent in cytokine-responsive cCREs in beta cells compared to alpha cells (∼15-25% beta cCREs, ∼5–15% alpha cCREs) (**Figure 2E, Table S12**). Similarly, a larger percentage of ER stress-responsive cCREs in beta cells had ATF4 and CHOP motifs compared to alpha cells (ATF4: 17.09% beta cCREs, 9.92% alpha cCREs; CHOP: 15.92% beta cCREs, 8.07% alpha cCREs) (**Figure 2E, Table S12**). Motif accessibility in individual cells supported stronger up-regulation of ATF and CHOP motifs in ER stress for beta cells (ATF4 beta cells avg. diff=2.119, alpha cells avg. diff=0.826) **(Figure 2F; Table S13**). Interestingly, RFX motifs showed opposed effects with decreased and increased accessibility in ER stress in beta and alpha cells, respectively (**Figure 2F; Table S13**). We defined TF gene regulatory networks (GRNs) by linking cREs to target genes using promoter proximity and Activity-By-Contact (ABC)^28^. For example, ATF4-regulated cCREs upregulated in beta cells under ER stress were linked to 32 genes with up-regulated expression, including ER stress regulators *ERN1, XBP1,* and *GADD45*^29^*, MAP3K5*^12^, and *GLIS3*^30,31^, as well as, interestingly, down-regulated genes such as *GCK*^32^ (**Figure 2G-H, Table S14**).

Finally, we determined to what extent genomic responses of islet cell types to ER stress were enriched in variants associated with T2D risk and fasting glucose (FG) level. We calculated gene p-values for T2D and fasting glucose (FG) level using MAGMA^33^ and tested whether genes up- and down-regulated in each stressor were enriched for T2D or FG association. We identified significant enrichment (FDR<0.01) of T2D and FG association among genes down-regulated in ER stress in beta cells (**Table S20, Figure 2I-J**), and this enrichment was highly specific compared to alpha cells as well as to inflammatory stress. Genes down-regulated in ER stress in beta cells further consisted of many involved in T2D and monogenic diabetes, including *SLC30A8* and *GCK* (**Table S8**). In addition, cCREs down-regulated in ER stress were significantly enriched (FDR=0.0016) for fine-mapped T2D variants (**Table S21**). Conversely, we found no significant enrichment of T2D variants among genes or cCREs up-regulated in inflammatory stress (**Table S21**).

Overall, these results demonstrate that beta cells have pronounced genomic responses to stress and that regulatory programs altered in beta cells in ER stress harbor T2D genetic risk.

### Network analysis links T2D genetic risk to beta cell molecular phenotypes and master regulators

We next sought to understand the effects of ER stress and inflammation on specific gene regulatory networks acting within beta cells during physiology and type 2 diabetes. We utilized multiome data generated in beta cells from un-stimulated islets^24^ to perform weighted gene co-expression network analysis (WGCNA). We identified 15 co-expression networks (M1-M15) in beta cells comprised of 3,412 genes **(Figure 3A, Table S15)**, which were highly reproducible across datasets (**Figure S4A**). Highly distinct pathways were enriched among the genes in each co-expression network such as regulation of inflammatory response for M4 (p=6.0×10^-4^), extracellular matrix organization for network M9 (p=0.004.45×10^-3^), chaperone-mediated protein folding for M12 (p=4.09×10^-5^), cellular lipid response for M2 (p=4.84×10^-9^), and core beta cell functions such as hormone secretion for M1 and M7 (p=1.95×10^-4^, p=1.71×10^-3^), revealing that networks reflect different biological processes acting within beta cells (**Table S16, Figure 3B, Figure S4B).**

**Figure 3.**
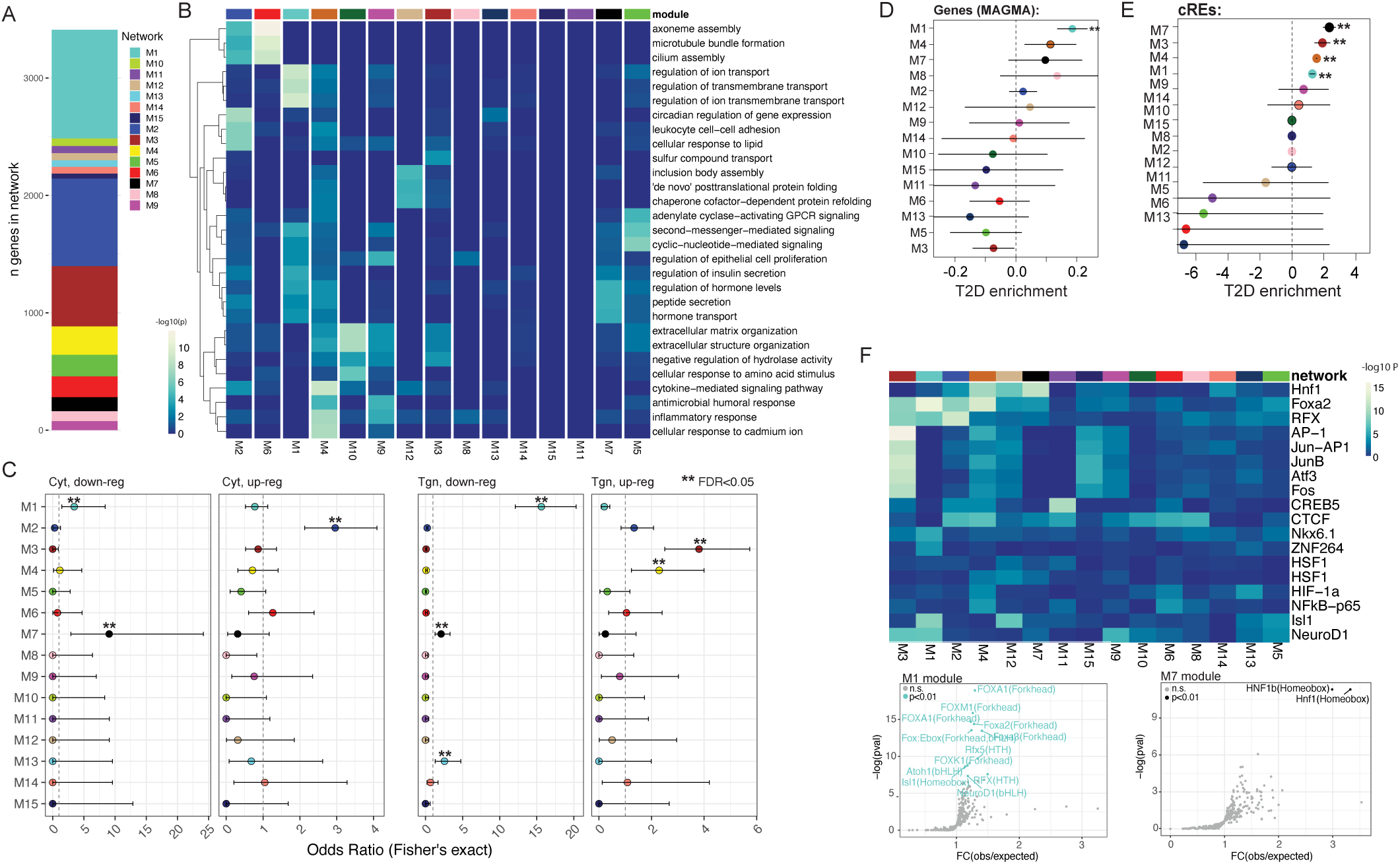
Gene co-expression networks in beta cells show altered expression in ER stress and enrichment for T2D risk. (A) Graphical representation of hdWGCNA networks. (B) Heatmap of selected biological processes enriched in hdWGCNA networks. P-values are from Goseq. (C) Fisher’s exact test for enrichment of cyt- and Tgn-beta cell differential genes with hdWGCNA networks. (D) Enrichment for T2D-associated genes in hdWGCNA networks. Enrichment scores are from MAGMA and based on summary statistics from Mahajan et al. (E) Enrichment of fine-mapped T2D risk variants in cCREs linked to genes in each network. Fine-mapping data from Mahajan et al. (F) Heatmap of enriched TF motifs in cCREs linked to gene networks (top) and volcano plots (bottom) highlighting the most enriched motifs in networks enriched for insulin secretion-associated genes. Key: **=FDR<0.05

We next determined the effects of stress on the activity of co-expression networks in beta cells. Genes in the insulin secretion-related networks (M1, M7) collectively showed significant down-regulation in stress (FDR<.05) and were enriched for genes down-regulated in ER and inflammatory stress (FDR<0.05). Down-regulated networks in ER stress were also enriched (FDR<.05) for marker genes of specific beta cell sub-types (beta-2 for M1, beta-1 for M7) with altered composition and secretory capacity in T2D^24^ (**Figure S4D, Table S19**). By comparison, the M3 and M4 networks associated with extracellular matrix organization and inflammation, respectively, had increased expression in ER stress (FDR<.05) and were enriched for genes upregulated by ER stress (FDR<0.05) (**Figure 3C, Table S18, Table S19, Figure S4C**). The T2D- and insulin secretion-enriched M1 network and the M3 network both had increased expression in T2D beta cells (FDR<0.05), highlighting signatures of beta cell ER stress altered in T2D (**Table S18, S19, Figure 4C**). The inverse activity of the insulin secretion-related M1 network in ER stress and T2D may, in the latter case, reflect increased insulin demand due to insulin resistance or medications that enhance insulin secretion.

**Figure 4.**
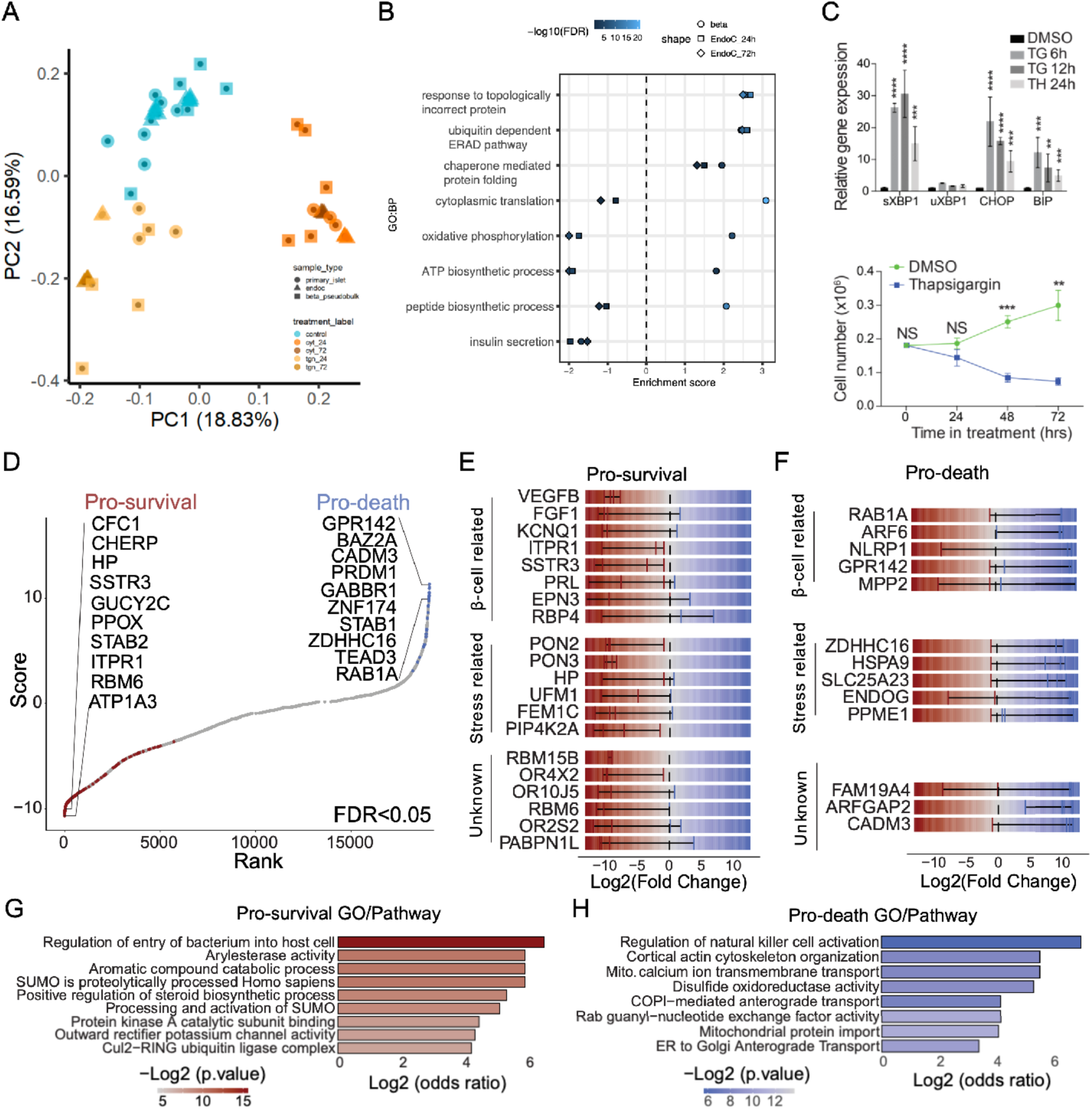
Genome-wide CRISPR screen identifies candidate genes influencing beta cell survival under ER stress. (A) Principal component analysis (PCA) of gene expression in bulk islets or pseudobulk beta cells treated or not with cytokines or thapsigargin for 24 hours, EndoC-βH1 cells treated or not with thapsigargin for 24 or 72 hours. (B) Normalized pathway enrichment scores (FGSEA) for selected GO:Biological Processses in pseudobulk beta cells or EndoC’s treated with Tgn for X hours. (C) qPCR results demonstrating activation of canonical UPR transcripts after 6, 12, or 24 hrs of Tgn exposure in EndoC-βH1 cells (top); Total number of EndoC-βH1 cells per 6-well plate after 0, 24, 48, or 72 hours of exposure to Tgn or an equivalent volume of DMSO (bottom). (D) Overview of candidate ER stress pro-survival and pro-death genes from pooled CRISPR-KO screen. (E) Line plot showing effect size per sgRNA for knockout of pro-survival CRISPR screen hits linked in previous literature to beta cells or stress responses, and CRISPR screen hits not previously associated with beta cells or stress responses. (F) Line plot showing effect size per sgRNA for knockout of pro-death CRISPR screen hits linked in previous literature to beta cells or stress responses, and CRISPR screen hits not previously associated with beta cells or stress responses. (G) Enriched gene ontology pathways in pro-survival genes from the CRISPR screen. (H) Enriched GO pathways in pro-death genes from the CRISPR screen.

Further underscoring the relevance of these co-expression networks to beta cell function and molecular phenotypes, M1 and M7 networks were significantly enriched (FDR<.05) for modulators of insulin content from a previous functional screen^14^ (**Figure S4D, Table S19**) and enriched (FDR<.05) for T2D and FG association, respectively (**Figure 4D, Table S20**). Intramodular connectivity analysis identified ‘hub’ genes in these networks with critical roles in beta cell function, including T2D risk gene *SLC30A8* in the M1 network and *HNF4A* in the M7 network (**Figure S4E, Table S15,17**). Additional T2D genes were highly connected to each network including *TCF7L2, ABCC8, RFX6,* and *GLIS3* in the M1 network and *HNF1A* and *SLC2A2* in the M7 network **(Figure S4E, Table S15,17**). The M4 network upregulated by ER stress (OR=6.61, FDR<0.05) was enriched for negative regulators of insulin content, highlighting the extensive genomic effects of ER stress on downregulation of insulin production.

We next sought to understand transcriptional drivers of co-expression network activity in beta cells. We used cCRE-gene links defined using promoter proximity and ABC to identify cCREs linked to genes in each network and conducted TF motif enrichment using the set of cCREs for each network. Overall, there were distinct sets of TFs enriched in cCREs linked to each network (**Table S17, Figure 3F, Figure S4F**). Motifs for known regulators of beta cell function such as FOXA2 and NKX6.1 were most enriched in T2D-enriched M1 network cREs, while HNF1 motifs were most enriched in fasting glycemia-associated M7 network cREs (**Figure 3F, Figure S4F, Table S17**), consistent with the motifs enriched in beta cell subtype-defining cCREs from our previous study^24^. By comparison, ATF and FOS/JUN motifs were most enriched in the M3 network which was up-regulated in ER stress. Fine-mapped T2D variants were significantly enriched (FDR<0.05) in cCREs linked to different networks both up-regulated (M1, M7) and down-regulated (M3, M4) by ER stress (**Figure 3E, Table S21**).

Overall, these results reveal gene co-expression networks in beta cells that are altered in ER stress, altered in T2D and affect genetic risk of T2D, as well as transcriptional regulators of these networks.

### Genome-wide screen identifies genes influencing beta cell survival in ER stress

Stress-induced genomic changes in beta cells may not directly translate to effects on cellular function, and we next sought to identify genes directly influencing beta cell function in ER stress. We utilized the beta cell line EndoC-βH1 to study the effects of genetic manipulations on cell survival under ER stress. The extent to which EndoC-βH1 cells effectively capture beta cell ER stress responses, however, has not been explored. We thus first performed RNA-seq on EndoC-βH1 cells treated with Tgn for 24 or 72 hours (n=3 each) and compared to baseline conditions. We conducted principal components analysis (PCA) of total RNA profiles from EndoC-βH1 cells, primary islets and primary beta cells with and without TGN or cytokine exposure. After accounting for donor effects, primary beta cells, primary islets and EndoC-βH1 cells subject to the same treatments broadly clustered together (**Figure 4A**), demonstrating high-level similarity in their transcriptomic profiles under ER stress.

Overall, gene and pathway expression changes showed a modest, positive correlation between EndoC-βH1 cells and primary beta cells treated with Tgn for 24 hours (r^2^=0.13, 0.24) and 72 hours (r^2^=0.13, 0.11) (**Tables S22-24**). Pathways related to UPR were highly upregulated across all sample types and time points, including up-regulation of master ER stress regulators *EIF2AK3*, *ATF6*, *HERPUD1*^34^, *HSP90B1*^34^ and the pro-apoptotic regulator *TMBIM6*^20^ (**Table S12, Table S22**) along with down-regulation of genes associated with insulin secretion such as *GCK* and *SLC30A8* (**Table S12, Table S22**). Gene and pathway expression profiles were overall highly correlated (r2=0.77) (**Table S23-24**) between EndoC-βH1 cells treated with Tgn for 24 or 72 hours. However, the magnitude of downregulation in peptide biosynthesis and chromatin organization pathways was greater at 72 hours, and certain pathways showed opposed effects at 24 and 72 hours of treatment, such as cell junction organization and chromatin organization (**Table S23-24**). Meanwhile, aerobic respiration was upregulated in primary beta cells and not in EndoC-βH1 cells (**Figure 4B**), revealing differences in how primary and immortalized cells respond to stress.

Next, we utilized this model to identify genes regulating beta cell survival and death under ER stress. We corroborated activation of canonical UPR following TGN treatment of EndoC-βH1 cells at 72-hour treatment using qPCR, as evidenced by altered splicing of *XBP1* and the upregulation of *BIP* and *CHOP* (**Figure 4C**). We next determined that 72-hour exposure to Tgn resulted in an approximate 30% survival rate in EndoC-βH1 cells (**Figure 4C**). We then performed a CRISPR/Cas9-mediated loss-of-function screen of survival in ER stress in EndoC-βH1 cells. We conducted the screen by transducing the human CRISPR Brunello sgRNA library into EndoC-βH1 cells. The representation of sgRNAs was evaluated 72 hours after exposing the transduced cells to either Tgn or control (DMSO) treatment. Overall, we identified 167 pro-survival genes and 47 pro-death genes in ER stress compared to control (FDR<.05) among 19,114 tested genes (**Table S25, Figure 4D-E**).

Pro-survival and -death genes included a subset with known roles in beta cell survival, function, or ER stress response. For example, among pro-survival genes, *SSTR3*^35^, *EPN3*^36^, and *PRL*^37^ (log2(FC)=-10.41, -4.70, -4.23) have been shown to protect beta cells against stress, while *ITPR1*^38^ and *FGF1*^39^ (log2(FC)=-10.48, -9.86) positively regulate beta cell mass under homeostasis and metabolic stress, respectively (**Table S25**). Notably, several negative regulators of beta cell insulin expression and secretion, including *KCNQ1*^40,41^, *VEGFB*^42,43^, and *RBP4*^44^ (log2(FC)=-10.039, -8.93; -4.39), were identified as pro-survival genes, highlighting an inverse relationship between insulin secretion and survival during ER stress (**Table S25**). For example, a *KCNQ1* mutation associated with neonatal diabetes has been shown to cause premature insulin hypersecretion and elevated apoptosis in stem cell-derived islet organoids^45^. Conversely, key mediators of stress-induced β-cell damage, such as the inflammasome factor *NLRP1*^46^ (log2(FC)=1.39) and the matrix metalloproteinase *MPP2*^47^ (log2(FC)=0.78), were identified among pro-death genes (**Table S25**). Several pro-death genes including *RAB1A*^48^, *GPR142*^49^, *ARF6*^50,51^, and *PARP2*^52^ (log2(FC)=10.15, 11.35, 9.89, 9.32) positively affect insulin secretion, again linking increased insulin secretion to decreased survival (**Figure 4E-F**).

Our screen also revealed genes not previously implicated in beta cells but that affect stress-induced cell death in other systems. For example, pro-survival genes *HP*^53^ and *PIP4K2A*^54^ (log2(FC)=-10.40, -8.61) are involved in preventing and resolving oxidative stress, and *PON2*^55^ and *PON3*^56^ (log2(FC)=-9.41, 9.27) (**Table S25**) protect cells from apoptosis during both oxidative and ER stress. Additionally, *FEM1C*^57^, and *UFM1*^58^ (log2(FC)=-8.79, -7.65) are associated with protein degradation and ER-phagy, critical mechanisms for mitigating ER stress (**Figure 4E-F, Table S25**). Supporting these findings, pro-survival genes were broadly enriched in pathways such as arylesterase activity (p=1.01×10^-3^, log2(OR)= 5.91) and the Cul2−RING ubiquitin ligase complex (p=6.78×10^-3^, log2(OR)= 4.21) (**Figure 4G, Table S26**). Pro-death genes mediated cell death through diverse mechanisms. For instance, *PPME1*^59^ (log2(FC)=1.21) is associated with oxidative stress, *ENDOG*^60^ and *ZDHHC16*^61,62^ (log2(FC)= 5.43, 10.227) are linked to DNA damage, *SLC25A23*^63^ (log2(FC)= 9.26) is implicated in mitochondrial calcium dysregulation, and *HSPA9* (log2(FC)=9.77) mediates calcium transfer from the ER to mitochondria, contributing to ER stress-induced oxidative stress and apoptosis^64^ (**Figure 4F**, **Figure 4H, Table S26**). Mitochondrial transmembrane transport (p=1.10×10^-3^, log2(OR)=5.54) and protein import (p=7.16×10^-3^, log2(OR)= 4.09), as well as Golgi-mediated transport (p=3.61×10^-3^, log2(OR)=3.40), were also broadly enriched among pro-death genes (**Figure 4G-H, Table S26**). Thus, altered ER and mitochondrial communication, as well as altered ER and Golgi communication, may promote beta cell death during ER stress. Finally, we identified regulators of beta cell survival with no prior known role in beta cell biology or stress responses such as RNA binding proteins (*RBM6*, *RBM15B*) (log2(FC)=-10.62, -8.99), cell-cell adhesion protein *CADM3* (log2(FC)= 10.94), GTPase-activating protein *ARFGAP2* (log2(FC)=9.94), and a novel cytokine ligand of formyl peptide receptor, *FAM19A4* (log2(FC)=5.44) (**Figure 4E-G, Table S26**).

Overall, these results reveal hundreds of genes mediating ER stress responses in beta cells which implicate novel biological processes in promoting cell survival and death in beta cells under ER stress.

### Beta cell survival genes under ER stress identify novel candidates for T2D risk

We determined the relationship between genomic changes and cell survival under ER stress. We tested genes with significant ER stress-responsive expression in beta cells or EndoC-βH1 cells for enrichment among genes affecting cell survival in ER stress. There was no significant enrichment (FDR>0.05) among these gene sets, including when splitting genes by up- or down-regulated expression or by pro-death or pro-survival (**Figure 5A, Table S27**). Similarly, pro-death and pro-survival genes in ER stress collectively showed no overall change in expression in beta cells in ER stress (NES=-0.797, p=0.83) (**Table S27**). Next, we tested genes linked to ER stress-responsive cCREs for enrichment among cell survival genes and similarly found no significant enrichment (FDR>0.05) (**Figure 5A, Table S27**). Finally, we examined enrichment among co-expression networks and again found no significant enrichment of genes in networks for cell survival genes (FDR>0.05) (**Table S27**). Overall, this suggests limited relationship between gene expression changes and functional effects on survival under ER stress in beta cells.

**Figure 5.**
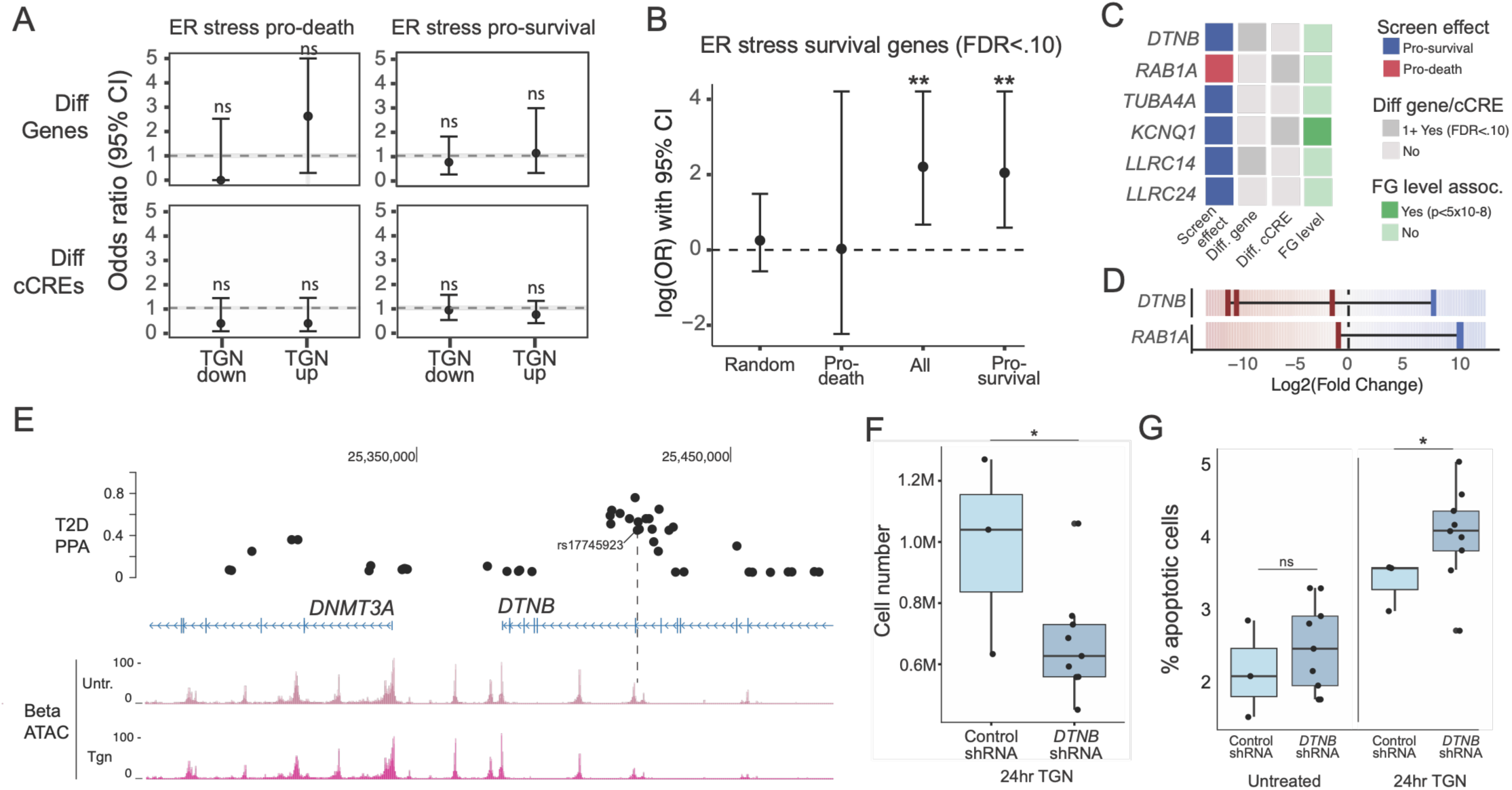
Genes promoting beta cell survival in ER stress affect T2D risk. (A) Enrichment of ER stress pro-death (left) and pro-survival (right) hits in differentially expressed genes (top) or target genes of differentially accessible cCREs (bottom) in beta cells exposed to Tgn. Effects are odds ratio and error bars are 95% confidence interval, and ns is not significant. (B) Enrichment of fine-mapped T2D risk variants in cCREs linked to pro-death genes, pro-survival genes, and a random set of 500 genes tested for differential expression in beta cells (random). Effects are odds ratio and error bars are 95% confidence interval, which are plotted on a log scale. (C) Summary of CRISPR screen hits linked to fine-mapped T2D risk variants by direction of screen effect (pro-survival, pro-death), whether gene expression was differential in beta cells under ER stress, whether 1+ cCRE linked to the gene had differential accessibility in beta cells under ER stress, and whether the gene is significantly associated (P<5×10^8^) with fasting glucose level. (D) Line plot showing effect size in fold-change per sgRNA on beta cell survival under ER stress for *DTNB* and *RAB1A.* (E) Genome browser tracks showing the *DTNB* locus with ATAC-seq signal in Tgn-treated and untreated beta cells, predicted cRE-gene links, and fine-mapping data for T2D. Fine-mapped variant rs17745923 overlaps a beta cell cRE in *DTNB*. (F) Quantification of cell number for EndoC-bH1 cells exposed to thapsigargin (TGN) treatment for 24hr in both non-targeting control shRNA (n=3) and *DTNB* shRNA (n=3 for each of 3 guides). P-values calculated using ANOVA, where * represents P<.05. Box plots represent median value with interquartile ranges. (G) Proportion of EndoC-bH1 cells from flow cytometry analysis with both FITC (ApoTracker) and propridium iodine (PI) markers in untreated and TGN treated conditions for 24hr in both non-targeting control shRNA (n=3) and *DTNB* shRNA (n=3 for each of 3 guides). P-values calculated using ANOVA, *P<.05 and ns=non-significant. Box plots represent median value with interquartile ranges.

Given limited evidence that cell survival genes have genomic changes in ER stress, we determined whether these genes showed evidence for a role in T2D genetic risk. We identified significant enrichment of T2D risk variants in cCREs linked to ER stress cell survival genes (log(OR)=2.05, p=6.0×10^-4^) (**Figure 5B, Table S28**), and this enrichment was more pronounced among genes promoting survival in ER stress (log(OR)=2.21, p=3.0×10^-4^) compared to pro-death genes (**Figure 5B, Table S28**). In contrast, cCREs predicted to regulate pro-survival genes under inflammatory stress showed no corresponding enrichment for T2D risk variants (log(OR)=-1.15, p=0.74) (**Table S28**). Further emphasizing the specificity of this enrichment, there was also no enrichment of T2D risk in cCREs linked to randomly selected genes with no effect on survival (log(OR)= 0.251, p=.26) (**Table S28**). Among cell survival genes linked to a cCRE harboring a fine-mapped T2D risk variant, almost all promoted survival under ER stress (**Figure 5C, Table S30**). As many T2D risk loci affect insulin secretion, we determined if cell survival genes were associated with measures of insulin secretion. Only one T2D-linked survival gene (*KCNQ1)* showed significant association with fasting glucose level, arguing that most are unlikely to affect T2D risk via strong effects on basal insulin secretion. Of note, multiple pro-survival genes (*DTNB, LRRC24, LRRC14)* were at loci in a previously defined cluster of ‘hyper insulin secretion’-related loci^65^, which is marked by compensatory insulin secretion.

We next highlighted specific pro-survival genes in ER stress linked to T2D risk variants. For example, *DTNB,* a structural protein involved in cytoskeletal organization, promoted beta cell survival in ER stress (**Figure 5C-D**). Beta cell cREs harboring fine-mapped T2D variants at this locus were linked to *DTNB* (**Figure 5E**). To validate the effects of *DTNB* on beta cell survival we performed shRNA-mediated knockdown in EndoC-βH1 cells under ER stress and assessed whether reduced expression of *DTNB* increased markers of apoptosis under untreated and ER stress conditions compared to a control transduced with non-specific shRNAs. We found a significantly reduced number of viable EndoC-βH1 cells and significantly higher levels of apoptosis in *DTNB* knockdown, which was specific to the ER stress treatment condition (ANOVA p-value<0.05; **Figure 5F-G**, **Figure S6**). Altogether, these findings support that *DTNB* activity promotes beta cell survival in ER stress and altered activity of this process may impact risk of T2D.

Overall, these results reveal T2D candidate genes that affect beta cell survival under ER stress including those not implicated by prior studies.

## Discussion

Genome-wide perturbation screens and single cell profiling uncovered novel mediators of ER stress responses in pancreatic beta cells and candidate genes for T2D. Several recent studies performed genomic profiling of islets under ER stress^12,13^ and annotated candidate genes at T2D loci based on genomic changes in stress. The results of our study highlighted many genes and pathways without prior known involvement in ER stress responses, and these genes did not broadly show changes in transcriptional levels in stress and were independently enriched for T2D loci. As T2D risk variants largely affect gene regulation, risk variants may alter the baseline expression of beta cell survival genes where the functional effects of altered expression then manifest under stress. Overall, survival genes represent a component of beta cell stress responses relevant to T2D risk not captured by previous studies that may represent novel targets for preserving beta cells in diabetes.

Pro-survival genes linked to T2D risk variants implicate diverse processes in T2D including cytoskeletal organization, signaling receptor activity, and ion channel activity. This includes *DTNB,* which encodes a component of the dystrophin-associated protein complex which links cytoskeleton to the extracellular matrix and has also been implicated in autism and Alzheimer’s disease^66,67,68^ and *TUBA4A* which is a component of microtubules previously linked to ALS^69,70,71^ and dementia^72^. Altered cytoskeletal activity may impact beta cell survival under stress in multiple ways including through regulation of ER stress sensors, interactions that drive remodeling of the ER, damaged connections within the cytoskeleton, and promotion of cell death and autophagy pathways. Pro-survival genes also did not collectively show effects on fasting glycemia, although several were implicated in hyper-insulin secretion^65^, and thus these genes may affect T2D risk in the context of increased insulin secretion demands during prolonged stress and T2D progression.

Our findings also underscore a relationship between insulin secretion and ER stress, where insulin secretion promotes ER stress and activation of the UPR, and prolonged insulin secretory demands can lead to beta cell dysfunction and death. Few genes known to play a role in insulin secretion showed effects on cell survival, suggesting that only a select subset of beta cell functions may lie at the intersection between the two processes. For example, *RAB1A* is implicated in vesicle trafficking^48,73,74^ which is likely relevant to both insulin secretion and cell survival under ER stress. Our results also highlight that therapies promoting insulin secretion may in some cases have detrimental effects on survival in ER stress; for example, ARF6 affects glucose-stimulated insulin secretion and is a potential therapeutic target^75,50,76^ but the results of our screen reveal that *ARF6* activity may also promote cell death under ER stress. Cell survival genes also provided mechanistic insights into pathways mediating ER stress responses beyond insulin secretion; for example, genes involved in mitochondrial transport pathways broadly promoted cell death in ER stress.

Our study also defined co-expression networks in beta cells that reflect genes involved in distinct biological processes operating within beta cells, as well as key transcriptional regulators of these networks. Several networks closely reflected previously described beta cell states which had distinct molecular profiles, exocytic properties under glucose stimulation, and abundance in T2D^77,24^. Our study revealed that these networks were also altered by ER and inflammatory stress, with stronger down-regulation of genes in the more highly exocytic state, mirroring the heightened exocytic defects observed in T2D donors^24^. Taken together, these observations reflect increased stress sensitivity in the more exocytic subset of beta cells, which may contribute to beta cell dysfunction in T2D progression as there are increased demands for insulin. Other networks were up regulated in ER stress but not altered in T2D and thus may reflect programs driving increased ER stress responses in beta cells in T2D progression prior to onset.

Limitations of the study include, first, that our functional screen compared the relative recovery of guides from cells between ER stress and normal conditions, and thus some genes identified by our screen may be related to cell proliferation instead of survival. Furthermore, our screen design does not enable identifying genes that may promote death or survival equally in baseline and ER stress conditions. Alternate screen designs, for example by sorting cells based on markers of ER stress or apoptosis, will help to overcome this limitation. In addition, some of the survival genes at T2D risk loci may function in other T2D-relevant cell types and therefore more systematic efforts to measure gene function across cell types are needed. Second, the screen was performed in the beta cell line EndoC-bH1, which is a model of primary beta cells. Although there were similar ER stress responses between EndoC-bH1 and primary beta cells, genome editing in primary cells or in stem cell-derived models may help reveal a more complete picture of genes affecting beta cell survival in ER stress. Finally, the use of a chemical model may mean that some responses differ from ER stress responses observed in physiology and T2D. Models of environmental contexts such as elevated glucose and lipid levels seen in glucolipotoxicity may help define changes that more closely mirror beta cell stress in T2D progression.

In summary, functional screens and single cell genomics provided novel insights into the genes and processes through which ER stress in beta cells contribute to T2D. These findings pave the way for mechanistic studies and therapeutic strategies to preserve beta cell function in T2D pathogenesis. More broadly, our findings highlight that a more comprehensive understanding of how cellular stress responses affect risk of disease requires functional read-outs beyond just genomic profiling.

## Acknowledgements

This work was supported by NIH awards DK122607 and HG012059 to K.J.G and M.S. This study includes data generated at the UC San Diego IGM Genomics Center utilizing an Illumina NovaSeq X Plus that was purchased with funding from a National Institutes of Health SIG grant (#S10 OD026929).

## Author contributions

M.O. performed genomics and gene function experiments, analyzed and visualized genomics and gene function data and wrote the manuscript. H.Z. performed gene function experiments and supervised the study. S.C., J.D, B.O. and K.V. performed genomics experiments. P.B. and R.E. analyzed genomic data. M.M. performed genomics experiments. A.W. supervised genomic data generation. M.S. obtained funding and supervised gene function experiments. K.J.G. obtained funding, supervised the study, analyzed data and wrote the manuscript.

## Supplemental tables

**Supplemental table 1.** Islet donor characteristics and summary of assays

**Supplemental table 2.** Differential gene expression results for cyt- and Tgn-treated bulk islets

**Supplemental table 3.** Differential chromatin accessibility results for cyt- and TGN-treated bulk islets

**Supplemental table 4.** Known motifs (HOMER) enriched in differentially accessible cCREs in bulk islets

**Supplemental table 5.** Enrichment for islet chromHMM states and differentially expressed genes in differentially accessible cCREs. Statistics reported are from the Fisher’s exact test. FDR was calculated separately for chromHMM tests and RNA-seq concordance, conducted post-hoc using the BH method.

**Supplemental table 6.** Gene ontology pathway enrichment for bulk DA-cCRE and DEG sets (GREAT binomial adjusted p-values and and Goseq over-representation p-value, respectively)

**Supplemental table 7.** Single cell library QC for multiome and snATAC-seq

**Supplemental table 8.** DESeq2 results for cyt- and TGN-treated pseudobulkbulk snRNA-seq

**Supplemental table 9.** Concordance of bulk islet differentially accessible cCREs and differentially expressed genes (DEGs) with and alpha cell and beta cell DA-cCREs and DEGs

**Supplemental table 10.** Pathway enrichment in cyt and tgn treated alpha and beta cells

**Supplemental table 11.** DESeq2 results for cyt and tgn treated snATAC-seq

**Supplemental table 12.** Enrichment of known motifs in alpha and beta cell DA-cCREs

**Supplemental table 13.** ChromVAR analysis in alpha and beta cells

**Supplemental table 14.** Gene regulatory network for ATF4

**Supplemental table 15.** Network assignments and gene connectivity scores

**Supplemental table 16.** Gene ontology enrichment in network gene sets

**Supplemental table 17.** Enrichment of known sequence motifs in network enhancer sets

**Supplemental table 18.** Differential network expression across conditions

**Supplemental table 19.** Enrichment for candidate regulators of insulin content or beta cell subtype marker genes in hdWGCNA networks

**Supplemental table 20.** MAGMA results

**Supplemental table 21.** Enrichment for fine-mapped T2D risk variants in sets of cCREs

**Supplemental table 22.** DESeq results for EndoC-βH1 cells

**Supplemental table 23.** Pathway enrichments EndoC-βH1 cells treated with cytokines or thapsigargin

**Supplemental table 24.** Correlation of DESeq test statistics in EndoC-βH1 and beta cells

**Supplemental table 25.** Gene-level effects on beta cell survival from CRISPR screen

**Supplemental table 26.** Gene set enrichment for beta cell survival genes

**Supplemental table 27.** Overlap between (DEGs) and genes linked to DA-cCREs and candidate regulators of beta cell survival

**Supplemental table 28.** T2D risk variant enrichment for CRISPR screen cRE sets

**Supplemental table 29.** Motifs with enrichment in cREs linked to pro-survival and pro-death genes

**Supplemental table 30.** Fine-mapped T2D risk variants linked to pro-survival CRISPR genes

**Supplemental Figure 1.**
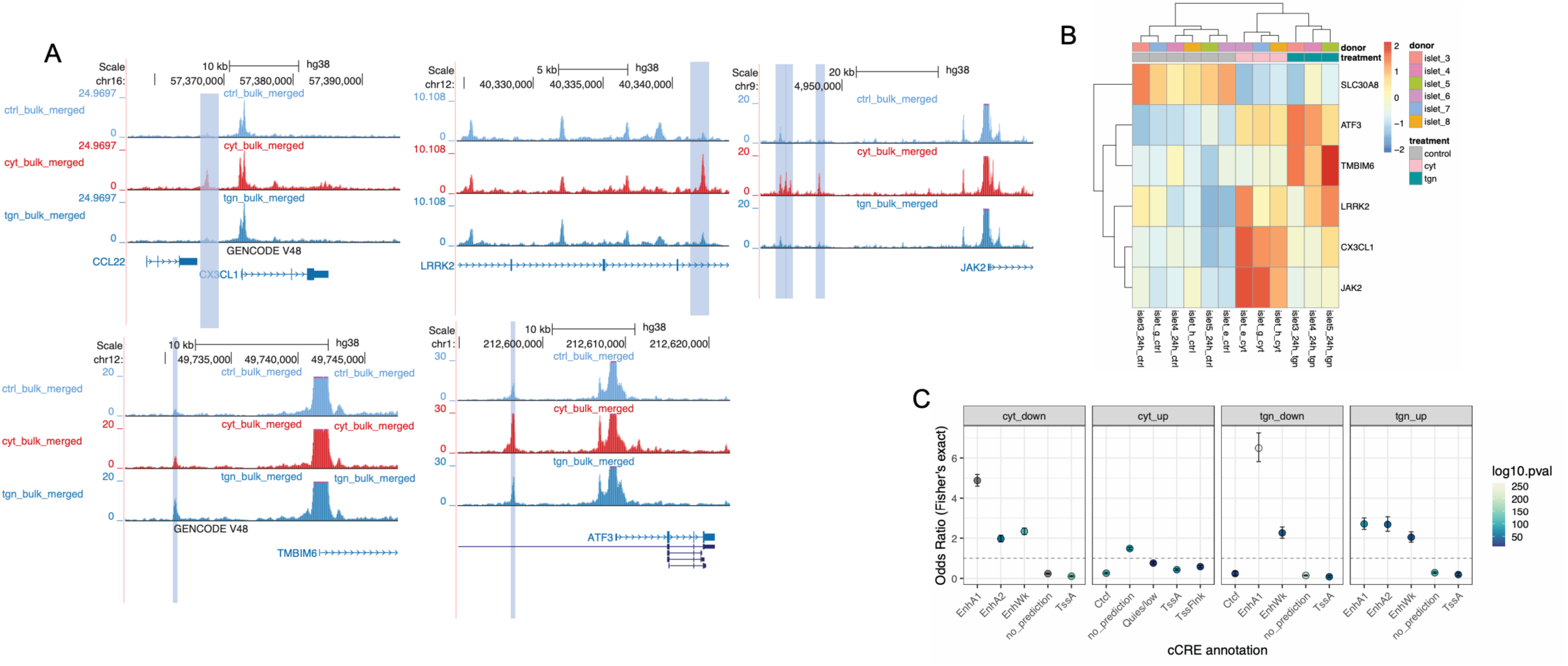
Exposure to pro-inflammatory cytokines or thapsigargin *in vitro* upregulate key pathways. (A) Heatmap of expression of selected genes with canonical roles in inflammation and ER stress. Counts normalized using the variance-stabilizing transformation from DESeq2 are displayed. (B) Genome browser tracks highlighting cCREs at or nearby differentially expressed genes with treatment-specific changes in accessibility. (C) Enrichment for cCREs overlapping islet chromatin states defined by chromHMM in a published study^22^ in differentially up- and down-regulated cCREs.

**Supplemental Figure 2.**
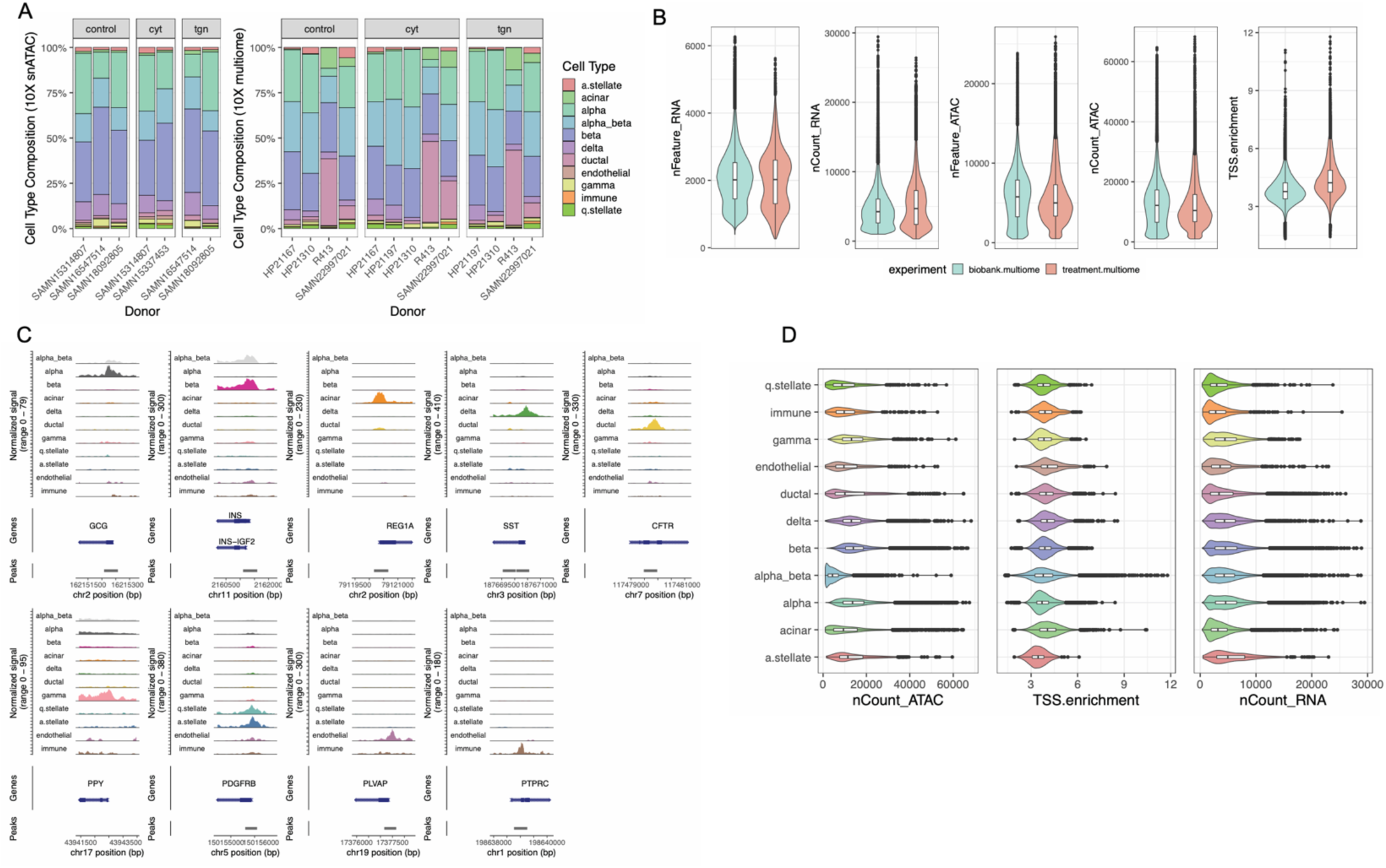
Single nucleus multi-omic profiling of chromatin accessibility and gene expression in islets under baseline, inflammatory, and ER stress conditions. (A) Overview of multiome and snATAC samples generated for this study showing cell type composition for each library. (B) Number of reads and features detected in multiome snRNA and snATAC, in addition to TSS enrichment in snATAC data. Our data (treatment.multiome) is shown separately from previously published islet multiome data (biobank.multiome) and is broadly comparable across measures of quality control. (C) Dot plot of marker gene expression for each cell type annotation. (D) Number of reads and features detected in the RNA and ATAC modalities in addition to ATAC TSS enrichment, split by cell type. Note low read counts in the snATAC modality for ‘alpha_beta’ cells

**Supplemental Figure 3.**
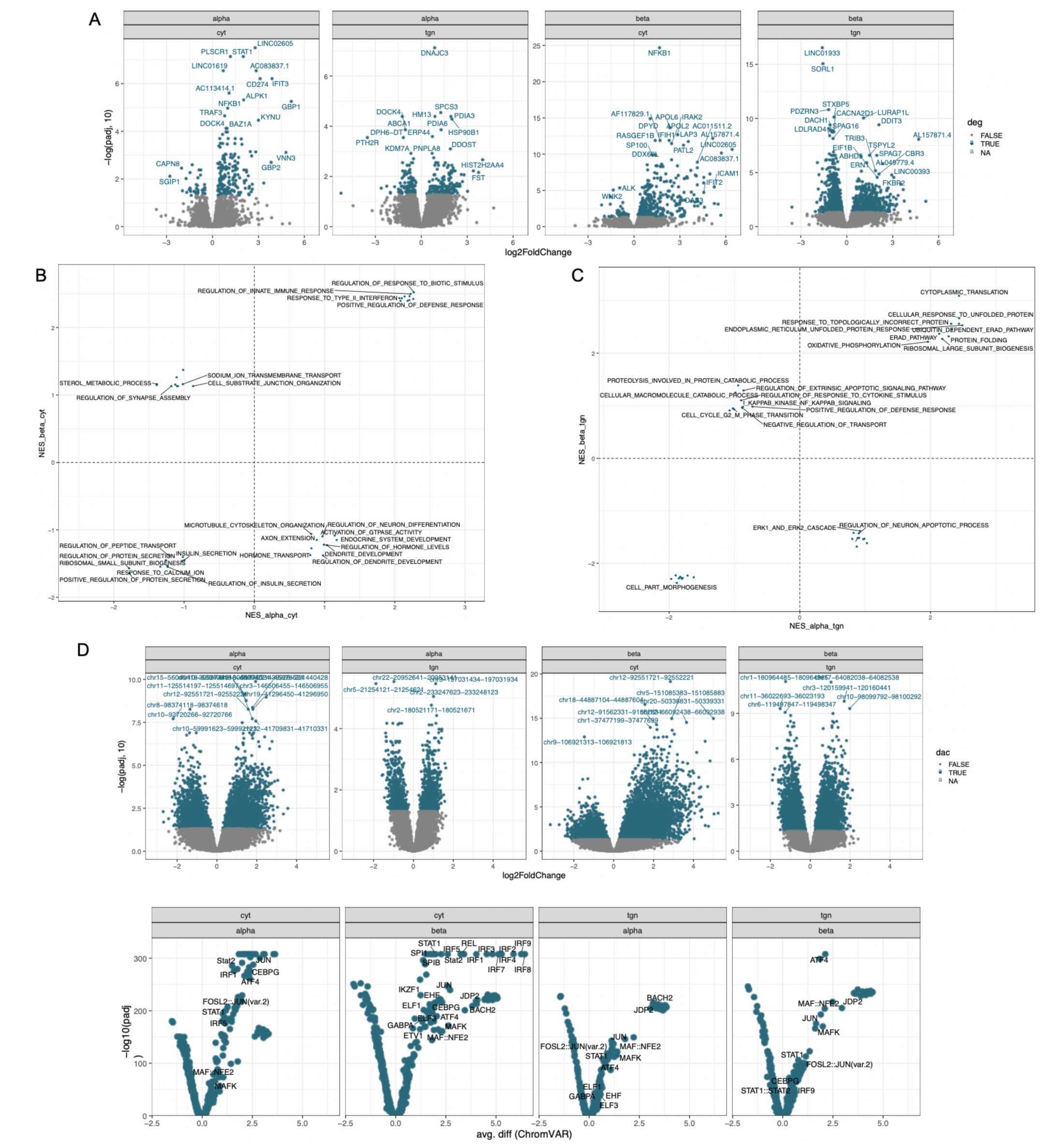
Cell type-resolved differential analysis of gene expression and chromatin accessibility under inflammatory and ER stress. (A) Counts for each cell type separated by treatment condition. (B) Volcano plots of differential expression and accessibility in alpha and beta cells under cyt and Tgn stimulation. (C) Volcano plots of motif activity (chromVAR) in alpha and beta cells under cyt and Tgn stimulation. (D) Heatmap of ChromVAR results in alpha and beta cells

**Supplemental Figure 4.**
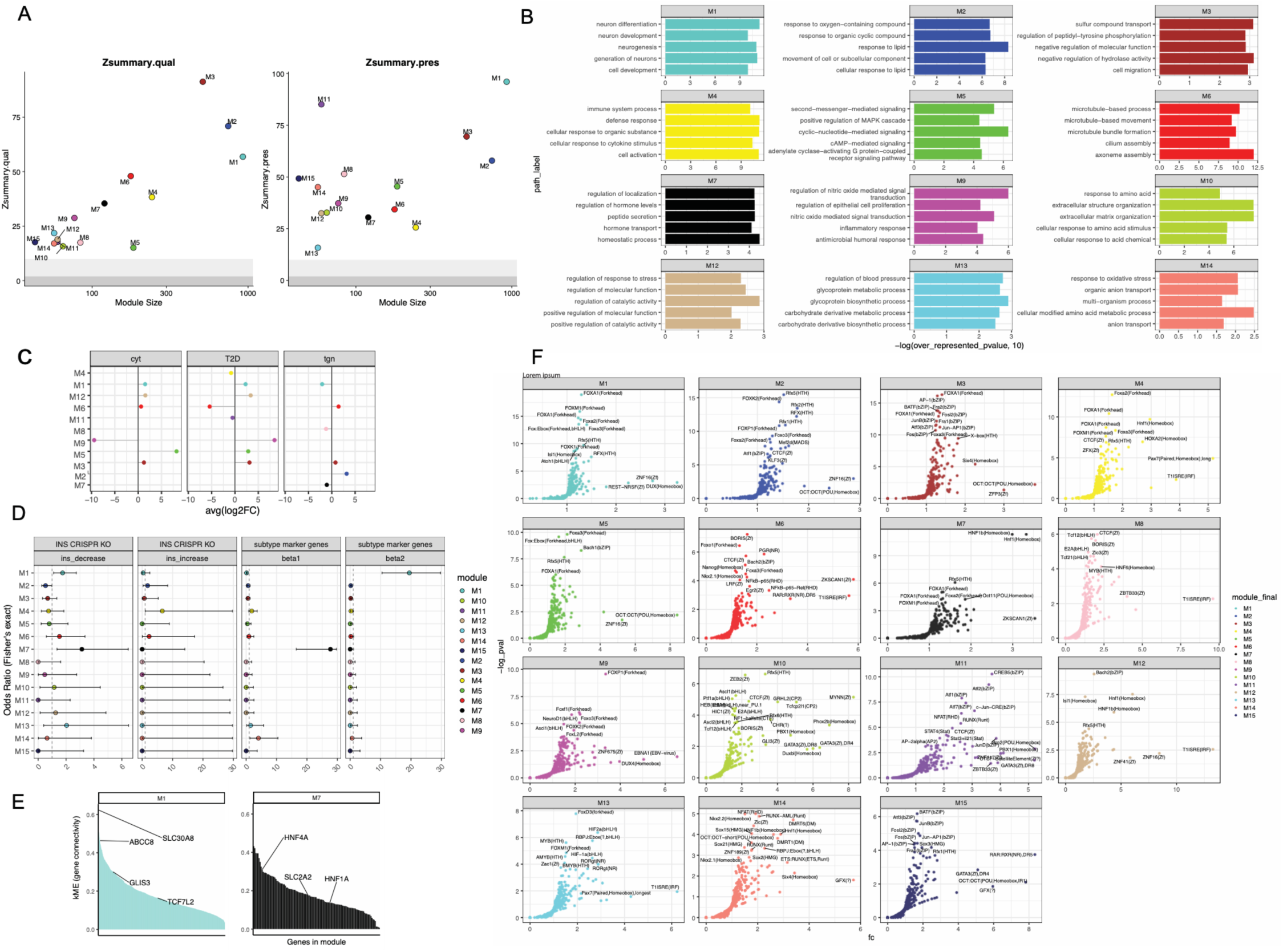
Annotation of co-expressed gene networks in unstimulated beta cells across T2D disease status. (A) Network reproducibility plots in our multiome dataset. (B) Enriched GO:BP pathways in networks. Over-represented p-values represented are from Goseq. (C) Differential network expression in cytokine- and thapsigargin-treated beta cells compared to unstimulated controls, and T2D beta cells compared to non-diabetic controls. Only significant contrasts (FDR<0.05) are displayed. (D) Fisher’s enrichment tests for overlap between gene networks and candidate regulators of insulin content identified via pooled CRISPR screen, or marker genes of functional beta cell subtypes. (E) Intramodular connectivity plots highlighting highly-connected ‘hub’ genes for networks enriched for association with T2D (top) or fasting glucose (bottom). (F) Motifs enriched in cCREs targeting genes in each network.

**Supplemental Figure 5.**
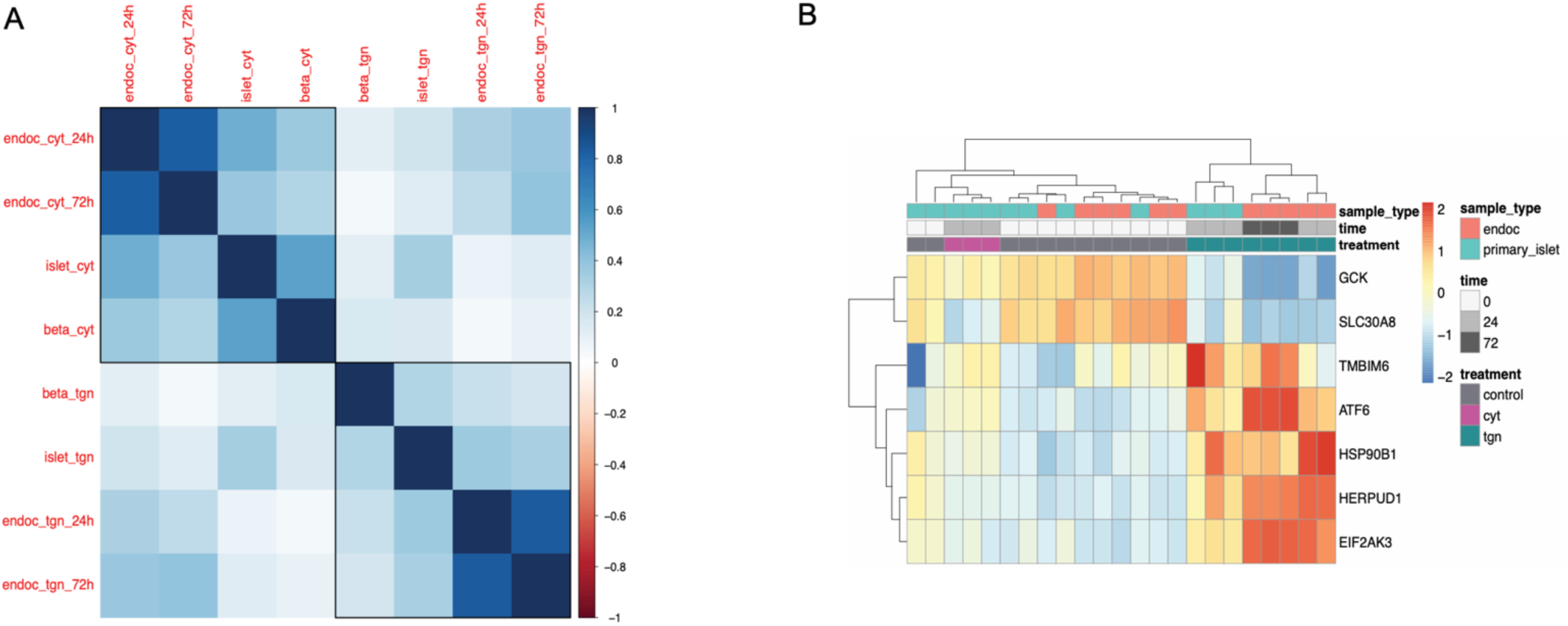
Evaluating an *in vitro* transcriptomic model of beta cell ER stress. (A) Transcriptome-wide correlation of cyt- and Tgn-induced gene expression changes in primary islets, pseudobulk primary beta cells, and EndoC-βH1 cells. (B) Normalized expression levels of selected genes in primary islets and pseudobulk beta cells, and EndoC-βH1 cells, treated or not with Tgn for the indicated number of hours.

## Methods

### Ethics statement

Pancreatic islet human donor tissue studies were approved by the IRB of the University of California San Deigo.

### *In vitro* treatments

To initiate treatments, the following stimuli were added to complete culture medium. For 72-hour treatments, media supplemented with treatment stimuli were refreshed daily. Thapsigargin (TGN) treatment: 1:1000 dilution of 1mM of thapsigargin (Thermo Fisher Scientific, T7459) dissolved in absolute DMSO. Cytokine (cyt) treatment: a cocktail of pro-inflammatory cytokines (10 ng/mL IFN-γ (Peprotech, 300-02), 0.5 ng/mL IL-1β (Peprotech, 200-01B), 1 ng/mL TNF-α (Peprotech, 300-01A)). For bulk RNA-seq and ATAC TGN treatments, controls were treated with an equivalent volume of DMSO. The effects of DMSO were found to be negligible based on PCA. All other ‘control’ samples were cultured in complete media only.

### Islet culture

Isolated islet preparations from non-diabetic donors were obtained from either the Integrated Islet Distribution Program (IIDP), Prodo Labs, and the Alberta Islet Core Distribution Program. Information on the islet donors is included in **Table S1**. We used a dithizone stain to hand-pick islets from non-endocrine debris. Islets were cultured overnight to recover from processing before initiating treatment conditions. Islets were cultured in CMRL1066 media (Invitrogen) supplemented with 10% FBS (Corning), 1X pen/strep (Invitrogen), 2mM L-glutamine (Cellgro), 1mM sodium pyruvate (Cellgro), 10µM HEPES (Gibco), and 0.25ug Amphotericin B, and a final concentration of 8mM D-glucose in 10cm dishes (Corning) at approximately 1-5 million islets per dish in a humidified incubator at 37C with 5% CO2. Islet studies were approved by the Institutional Review Board of the University of California San Diego.

### Bulk islet ATAC-seq data generation

We conduct ATAC-seq as described previously^79^. Cultured islets were resuspended in a chilled nuclei permeabilization buffer composed of 5% bovine serum albumin (BSA), 0.2% IGEPAL CA-630, 1 mM dithiothreitol (DTT), and 1X cOmplete EDTA-free protease inhibitor cocktail (Sigma-Aldrich) in 1X phosphate-buffered saline (PBS). The suspension was homogenized 10X with a “tight” pestle in a chilled dounce homogenizer and subsequently incubated on a tube rotator for 10 minutes to facilitate permeabilization. The homogenate was then passed through a 30 µm filter (Sysmex) to remove debris and centrifuged at 500 xg for 5 minutes at 4C to pellet the nuclei. The resulting nuclear pellet was resuspended in 1X TDE1 buffer (Illumina) and nuclei were counted using the Countess II Automated Cell Counter (Thermo Fisher Scientific). Approximately 50,000 permeabilized nuclei per sample were tagmented in a 25 µL reaction containing Tagmentation buffer and 2.5 µL of TDE1 enzyme (Illumina). The mixture was gently pipetted to mix and incubated at 37°C for 30 minutes in a thermal cycler to facilitate transposition. Following the reaction, DNA was purified using a 2:1 volume of AMPure XP beads (Beckman Coulter) and eluted in 20 µL of Buffer EB (Qiagen).

Library amplification was carried out using 10 µL of tagmented DNA in a 25 µL PCR reaction, prepared with the Nextera XT dual-index primer set (Illumina) and NEBNext High-Fidelity PCR Master Mix (New England Biolabs). The PCR program included an initial extension at 72C for 5 minutes, followed by denaturation at 98C for 30 seconds, then 12 cycles of 98C for 10 seconds and 63C for 30 seconds, with a final extension at 72C for 1 minute.

Size selection was performed in two steps using AMPure XP beads: first, 0.55X the library volume of beads was added and incubated for 15 minutes; the supernatant was transferred to a new tube, and an additional 0.65X volume of beads was added and incubated for 15 minutes. Libraries were eluted in 20 µL of Buffer EB.

Library quality and concentration were assessed using the Qubit High Sensitivity DNA assay (Thermo Fisher) and a 2200 Bioanalyzer (Agilent). Libraries were sequenced at the UC San Diego Institute for Genomic Medicine on an Illumina HiSeq 4000 platform with 100 bp paired-end reads, generating an average of 50 million read pairs per sample.

### Bulk islet ATAC-seq processing

Adapter sequences are trimmed from reads using TrimGalore^80^ with flags ‘–paired’ and ‘–quality 10’. Then, the trimmed reads are aligned to human genome build hg38 with BWA with the ‘-M’ flag and duplicate reads are removed using Picard MarkDuplicates^81^. We call peaks from de-duplicated reads for each sample with MACS2^82^ with parameters ‘–extsize 200 –keep-dup all –shift -100 –nomodel’.

### Bulk islet ATAC-seq count matrix generation PCA

To create a “master” set of cCREs profiled across all treatments, we collapsed peaks called in each sample into a master list by signal using BEDOPS. We then generated read counts at each cCRE in the “master” list for each sample using FeatureCounts^83^. The resulting read count matrix was transformed using the variance-stabilizing transformation from DESeq2^84^, and limma was used to regress out batch effects. Principal components were calculated using the ‘prcomp’ function. The top two principal components were then plotted with lines connecting treated samples to the respective donor-matched untreated control.

### Bulk RNA data generation

Total RNA was isolated using the RNEasy Mini kit (Qiagen). Approximately 500 IEQs per sample were used for bulk islet RNA-seq. These samples were lysed by vortexing at maximum speed in 350µL Buffer RLT (Qiagen). For RNA-seq experiments in EndoC-βH1 cells, 1 well of a 6-well plate of EndoC per sample was lysed by adding 350µL of buffer RLT directly to each sample, then scraping samples and transferring lysates to a 1.5mL Eppendorf tube before vortexing at maximum speed. RNA concentration was measured using a Nanodrop D1000 and adequate RNA integrity was verified using a 2200 TapeStation (Agilent).

Ribosomal RNA-depleted, strand-specific RNA-seq libraries were constructed by the UCSD Institute for Genomic Medicine using the TruSeq Stranded Total RNA Library Prep Gold kit (Illumina, Cat# 20020599). Libraries were sequenced at the UCSD Institute for Genomic Medicine on an Illumina HiSeq 4000 platform using 100 bp paired-end reads and targeting 50 million reads per sample.

### Bulk islet RNA-seq data processing

We used salmon^85^ with the flags ‘–validateMappings’ ‘—gcBias’ to quantify expression of human transcripts in Gencode v41.

### Bulk islet RNA-seq count matrix generation and PCA

We used the tximport package to import transcript quantification estimates from salmon^85^ to a read count-by-sample matrix. For principal component analysis, we used the variant-stabilizing transformation and limma to regress out ‘batch’ and ‘donor’ effects. Principal components analysis was performed using the ‘prcomp’ function without scaling. The top two principal components were then plotted with lines connecting treated samples to their donor-matched untreated control.

### Differential analyses in bulk islets

Read count matrices for bulk islet ATAC-seq and RNA-seq as described above were used for differential gene expression or cCRE accessibility analysis. We used the experimental design ‘∼treatment + donor’ to enable comparison of treatment effects within donor. We used the Wald test with a local fit dispersion for all differential analyses. We considered cCREs or genes significant at padj<0.05.

### Enrichment of DA-cCREs in proximity to DEGs

We used the Signac function ClosestFeature() to assign cCREs to a target gene if it is within 10kb of the closest gene feature and calculated enrichment using a Fisher’s exact test, using the set of all genes that passed DESeq2’s filtering cutoffs to define the “universe” of genes. We used the Bonferroni method to adjust p-values based on the total number of tests performed.

### Enrichment of chromatin states in DA-cCREs

We used liftOver to transfer islet-specific chromatin state predictions from a published study15 from hg19 to hg38. Then, we used bedtools closest to identify the closest state prediction to each cCRE. If the closest state prediction was >500bp from a cCRE, we labelled the state of the cCRE as “no_prediction”. If multiple chromatin states overlapped a cCRE, we labelled the cCRE with both annotations. We then used a Fisher’s exact test to quantify enrichment for each chromatin state in sets of DA-cCREs and used the Bonferroni method to adjust p-values based on the total number of tests performed.

### Bulk RNA-seq pathway enrichment

Sets of genes up- or down-regulated in each treatment condition were input to GOSeq. The set of all genes passing DESeq2’s filtering cutoffs in either the cyt vs. control or TGN vs. control contrasts were used as background.

### Bulk ATAC-seq pathway enrichment

Sets of cCREs up- or down-regulated in each treatment condition were input to rGREAT^86^ specifying the hg38 genome and otherwise default settings.

To arrive at the set of pathways displayed in Figure 1, we took the union of the top 10 pathways from the Gene Ontology biological process database that were enriched in each gene or cCRE set as ranked by adjusted p-value. Clustering of pathways by p-value across all DA-cCRE and DEG sets was calculated by pheatmap using Euclidean distance. We divided clusters into 3 sections with the pheatmap function cuttree_rows for aesthetic purposes only.

### HOMER motif analysis

Sets of cCREs were analyzed for TF motif enrichment compared to a background of the union set of cCRE regions in bulk islets or snATAC-seq, respectively, using HOMER’s findMotifsGenome.pl function with the command “perl findMotifsGenome.pl [.bed file of sites] hg38 [output directory] -bg [.bed file of background sites] -size 200 -p 20 -bits”.

### EndoC-βH1 culture

EndoC-βH1 cells between passages 39-60 were maintained according to established culture methods. Cells were grown in vessels pre-coated with ECM (Sigma, E1270) and Fibronectin (Sigma, F1141) supplemented with 1% Penicillin-Streptomycin (Gibco, 15140122). Complete cell culture media containing low glucose DMEM (Gibco,11885084), 2% BSA (Sigma, A1470), 50µM 2-mercaptoethanol (Gibco, 21985023), 0.12% Nicotinamide (Calbiochem, 481907), 5.5 ng/mL transferrin (Sigma, T8158), 6.7 pg/mL Sodium Selenite (Sigma, 214485) and 1% Penicillin-Streptomycin were refreshed every 2 days. Cells were passaged weekly using 0.25% Trypsin-EDTA for dissociation, which was quenched with an equal volume of heat inactivated FBS and two volumes of IMDM media (Gibco, 12440053). Dissociated cells were spun down at 500 rcf for 5 min at room temperature and counted before passaging cells into fresh, pre-coated culture vessels. Cells are tested at least every 10 passages to confirm the absence of mycoplasma contamination.

### EndoC RNA-seq data generation

RNA-seq assays, n=3 technical replicates were collected for each treatment at the indicated time points (t=24h, 72h). n=3 untreated control samples from the same passage and cultured in an identical format were collected alongside each set of samples. Each sample comprised of approximately 1 million EndoC-βH1 cells passaged from continuous culture and cultured a 6-well plate overnight before initiating treatments. Samples were lysed by adding 350µL of buffer RLT directly to each well and samples were scraped to facilitate lysis. Lysates were transferred to a 1.5mL Eppendorf tube and vortexed at maximum speed. RNA concentration was measured using a Nanodrop D1000 and adequate RNA integrity was verified using a 2200 TapeStation (Agilent).

Ribosomal RNA-depleted, strand-specific RNA-seq libraries were constructed by the UCSD Institute for Genomic Medicine using the TruSeq Stranded Total RNA Library Prep Gold kit (Illumina, Cat# 20020599). Libraries were sequenced at the UCSD Institute for Genomic Medicine on an Illumina HiSeq 4000 platform using 100 bp paired-end reads and targeting 50 million reads per sample.

### EndoC RNA-seq count matrix generation and PCA with primary islet RNA-seq and pseudobulk beta cell snRNA-seq

We used the tximport package to import transcript quantification estimates from Salmon to a read count-by-sample matrix. For principal components analysis, the EndoC RNA-seq count matrix was combined with the bulk islet RNA-seq and pseudobulk beta cell count matrices and normalized with the the variant-stabilizing transformation. We use limma to regress out ‘batch’ and ‘donor’ effects, treating all EndoC samples as the same ‘donor’ and samples collected at each time point as separate batches. Principal components analysis was performed on this normalized, batch- and donor-corrected TPM matrix using the ‘prcomp’ function without scaling. The top two principal components were plotted using ggplot2 ‘autoplot’ function.

### Differential expression analysis in EndoC-βH1

Transcript quantification estimates from salmon were used for differential gene expression analysis using DESeq2. We used the design ‘∼time_point + treatment’ to compare gene expression in cyt- or TGN-treated samples to controls collected at the same time point. We considered genes significant at padj<0.05.

### CRISPR screen

Genome-wide CRISPR screen was conducted as described in our previous publication. In brief,the EndoC-βH1 cells were expanded to a total of 300 million cells before spin-inoculation with the lentiviral human CRISPR Brunello library at an MOI=0.1. To enrich for successfully transduced cells, a 3-day puromycin (5 μg/mL, Sigma, P8833) selection was performed 48 hours after the spin-inoculation. Approximately 30M (400X genome coverage) cells were harvested as a representation control for the Brunello sgRNA library. The rest of the cells were kept in culture for an additional 14 days with puromycin (1 μg/mL) to achieve sufficient gene deletion and were subsequently treated with either 1 μM thapsigargin or an equal volume of DMSO for 72 hours. A time-point experiment was performed to evaluate which treatment duration was necessary to induce cell death. EndoC-βH1 cells were seeded 24 hours before the thapsigargin treatment and residual cell number was counted at 0, 24, 48 and 72 hours of treatment (n=3). Additionally, mRNA expression level of gene markers of ER stress is evaluated through RT-qPCR at 6, 12 and 24 hours of treatment (n=3). We harvested another 30M (400X genome coverage) cells from the DMSO treated cells and 15M (200X genome coverage) cells from the thapsigargin treated cells, although they started with the same number.

Genomic DNA from all three conditions were purified with a Quick-gDNA™ MidiPrep kit (Zymo Research, D3100). gRNA libraries were amplified from the genomic DNA using a two-step nested PCR. In brief, guide RNA inserts were amplified from the genomic DNA with the following primers:

F1-1:TCCCTACACGACGCTCTTCCGATCTNNNNNGGAAAGGACGAAACACCG

F1-2:TCCCTACACGACGCTCTTCCGATCTNNNNNHGGAAAGGACGAAACACCG

F1-3:TCCCTACACGACGCTCTTCCGATCTNNNNNHHGGAAAGGACGAAACACCG

F1-4:TCCCTACACGACGCTCTTCCGATCTNNNNNHHYGGAAAGGACGAAACACCG

R1-1:GGAGTTCAGACGTGTGCTCTTCCGATCNNNNNTGCTATTTCTAGCTCTAAAAC

R1-2:GGAGTTCAGACGTGTGCTCTTCCGATCNNNNNVTGCTATTTCTAGCTCTAAAAC

R1-3:GGAGTTCAGACGTGTGCTCTTCCGATCNNNNNVMTGCTATTTCTAGCTCTAAAAC

R1-4:GGAGTTCAGACGTGTGCTCTTCCGATCNNNNNVMAATGCTATTTCTAGCTCTAAAAC. Pooled F1 primers (F1-1 to F1-4) and R1 primers (R1-1 to R1-4) were used in each PCR reaction to avoid cluster registration failure on Illumina machines. Amplicons from the first step of PCR were gel purified and subjected to a second round of PCR to add Illumina sequencing adaptors and TruSeq indexes. Primers used in the second PCR step are the following: F2:AATGATACGGCGACCACCGAGATCTACACTCTTTCCCTACACGACGCTCTTCCGA; R2:CAAGCAGAAGACGGCATACGAGATNNNNNNGTGACTGGAGTTCAGACGTGTGCTCTTCCG.

Sequencing library amplified from the second round of PCR were size-selected and purified with a magnetic bead-based SPRIselect reagent (Beckman Coulter, B23318), and subjected to Nova-seq 6000 Illumina NGS platform using a pair-end sequencing (PE150) method at the UCSD Institute for Genomic Medicine.

### Analysis of CRISPR screen results

A UMI identification method was incorporated into the pipeline by counting species of variable nucleotides from F1 and R1 primers above. Following UMI identification, adaptor sequences ggaaaggacgaaacaccg and gttttagagctagaaatagca flanking the 19-20 base pair of sgRNA sequences were trimmed using cutadapt. Trimmed sequencing reads were then aligned to the reference sgRNA library with bowtie2 with default settings^87^. The output BAM files were subjected to the ‘umi_tools’ pipeline^88^ for removal of sequence duplication, resulting a de-duplicated BAM file that can be used for sgRNA counting with the MAGeCK model-based tool for CRISPR-Cas9 knockout screens. Statistical significance of guide RNA representation in DMSO and thapsigargin treated datasets for n=19,114 genes was estimated with the ‘rra’ subcommand in the MAGeCK package and p-values were corrected using FDR. Genes with significantly enriched (47, pro-death) or depleted (167, pro-survival) sgRNAs in the thapsigargin treated group at FDR<.05 were extracted as top candidate regulators.

### Gene knockdown experiments

One day prior to transfection, approximately 500,000 viable EndoC-βH1 cells/well were seeded into 12-well plates. Cells were spin-transduced at 980xg for 1 hour with either commercially prepared lentivirus particles containing shRNAs specific to the *DTNB* gene, or lentiviral particles containing non-targeting scramble shRNAs, in the presence of 8ug/mL of polybrene. Transfection media was replaced by complete culture media the next morning and cells were cultured for 4 days, with media replaced every other day, to recover from transduction. On day 5 following transduction, indicated samples were treated with 1µM of thapsiargin for 24 hours. Following thapsigargin treatment, samples were trypsinized according to standard protocol and resuspended in 500µL of 0.2% DPBS/BSA-V. Cell counts and viability were determined using a Countess 2 cell counter with Trypan blue staining (Invitrogen).

To assess apoptosis, we stained cells transduced and counted as described above using 80µM ApoTracker Green (BioLegend) in 0.2% DPBS/BSA-V. Samples were subsequently washed twice with 0.2% DPBS/BSA-V and resuspended in 0.2% DPBS/BSA-V containing 10% propidium iodide (PI). Samples assessed with flow cytometry on a BDSLR Fortessa-X20. Flow cytometry data was analyzed using FlowJo 10.

We tested for differences in cell numbers and apoptotic cell proportion (FITC+/PI+) between *DTNB* shRNA guides and non-targeting control shRNA in each treatment using two-way ANOVA.

### Isolation of islet nuclei for 10X single-nucleus assays

Approximately 1,000 frozen islet equivalents (°1x 10⁶ cells) were pulverized using a mortar and pestle on liquid nitrogen. Ground tissue was resuspended in 1 mL of nuclei-permeabilization buffer composed of 10 mM Tris-HCl, pH 7.5, 10 mM NaCl, 3 mM MgCl₂, 0.1 % Tween-20 (Sigma), 0.1 , IGEPAL-CA630 (Sigma), 0.01 % digitonin (Promega) and 1 % fatty-acid-free BSA (Proliant, 68700) in molecular biology-grade water followed by dounce homgenization (Wheaton) with a tight pestle for 10–20 strokes to obtain a uniform nuclei suspension. Homogenized islet suspension was passed through a 30-µm cell strainer (Sysmex) and then rotated at 4 °C for 10 min to before they were collected by centrifugation at 500 × g for 5 min (fixed-angle rotor, 4 °C).

### Single-modality 10X snATAC-seq data generation

Permeabilized islet nuclei, prepared as described above, were washed once with wash buffer containing 10mMTris-HCl, 10mM NaCl, 3mM MgCl₂, 0.1% Tween-20, and 1 % BSA in molecular biology-grade water and resuspended in 30 μl of 1x Nuclei Buffer (10x Genomics). Nuclei were counted using a hematocytometer and approximately 15,360 nuclei per sample were used for 10X snATAC-seq. Libraries were generated with the Chromium Single Cell ATAC Library & Gel Bead Kit (10× Genomics, Cat #1000110) together with the Chromium Chip E (Cat #1000086) and Chromium i7 Multiplex Kit (Set A, Cat #1000084), following the manufacturer’s protocol. Final libraries were quantified on a Qubit fluorometer (Life Technologies) and fragment size distribution was inspected on an Agilent TapeStation (High-Sensitivity D1000) to confirm the characteristic nucleosomal ladder indicative of successful tagmentation. Libraries were sequenced on an Illumina NovaSeq 6000 using a 50 + 8 + 16 + 50 bp read configuration (Read 1, i7 index, i5 index, Read 2).

### 10X snMultiome-seq library generation and sequencing

Nuclei were resuspended in 400 µL of sort buffer consisting of 1X protease inhibitor cocktail (Thermo Fisher, 1 U/µL RNasin (Promega) and 1% fatty acid-free BSA in 1X PBS and stained with 1 µM 7-AAD (Thermo Fisher, A1310). Approximately 120,000 nuclei were sorted on a Sony SH800 into 87.5 µL of collection buffer (1 U/µL RNasin, 5 % fatty acid-free BSA in 1X PBS). Sorted nuclei were mixed 4:1 with 5× permeabilization buffer, containing 50 mM Tris-HCl pH 7.4, 50 mM NaCl, 15 mM MgCl₂, 0.5 % Tween-20, 0.5 % IGEPAL-CA630, 0.05 % digitonin, 5 % BSA, 5 mM DTT, 5× protease inhibitors, and 1 U/µL RNasin in molecular biology-grade water. After a 1-min incubation on ice, nuclei were pelleted at 500 × g for 5 min in a swinging-bucket rotor at 4C. Supernatant was carefully aspirated, retaining approximately 50 µL of supernatant to minimize loss of pelleted nuclei. 650 µL of wash buffer (10 mM Tris-HCl pH 7.4, 10 mM NaCl, 3 mM MgCl₂, 0.1 % Tween-20, 1 % BSA, 1 mM DTT, 1× protease inhibitors, 1 U/µL RNasin in molecular biology-grade water) was added gently and the suspension was centrifuged again under the same conditions before the supernatant was removed, leaving ∼2–3 µL with the pellet. 7–10 µL of 1× Nuclei Buffer (10× Genomics) was used to gently resuspend nuclei, and resuspended nuclei were counted on a hemocytometer. Approximately 16,550–18,000 nuclei per sample were used for multiome library generation; for pooled samples, an equal number of nuclei from each sample in the pool were combined after nuclei quantification and preceding GEM formation. Multiome libraries were prepared with the Chromium Next GEM Single Cell Multiome ATAC + Gene Expression Reagent Bundle (10× Genomics, Cat #1000283) and accompanying chip (Cat #1000234) per the manufacturer’s instructions, together with Dual Index Kit TT Set A (Cat #1000215) and Single Index Kit N Set A (Cat #1000212).

Libraries were amplified under the following PCR cycling conditions: 7 cycles of ATAC index PCR; 7 cycles of cDNA amplification; 13-16 cycles of RNA index PCR depending on cDNA yield. Resulting libraries were quantified on a Qubit and ATAC library fragment sizes were assessed on an Agilent TapeStation (High-Sensitivity D1000). Libraries were sequenced on a NovaSeq 6000 S4 run using the following configurations: ATAC: 50 + 8 + 24 + 50 bp (Read 1, i7, i5, Read 2); RNA: 28 + 10 + 10 + 90 bp (Read 1, i7, i5, Read 2).

### Quality control and clustering of snATAC and snMultiome-seq

Multimodal snATAC+RNA data was aligned to hg38 using 10X Genomics CellRanger-ARC v2.0.1, while single-modality snATAC data was aligned to hg38 with 10X Genomics CellRanger v6.0.1. Several per-cell quality control metrics are used to identify high-quality cells for downstream analysis. For the RNA modality, we retained cells with a log10 RNA read count between 3.5 and the top 1% of each dataset, more than 1,500 detected genes, and less than 15% mitochondrial transcripts. For the ATAC modality, we required a log10 ATAC read count between 3 and the top 1%, 2,000 to 20,000 fragments in peak regions, more than 15% of reads in called peaks, less than 5% of reads mapping to blacklist regions, a nucleosome signal below 4, and a TSS enrichment score above 2.

Additionally, we corrected for ambient RNA contamination using SoupX’s autoEstCont method^25^ for each sample and used Scrublet^89^ and AMULET^26^ with default settings to identify and remove doublet cells from the RNA-seq and ATAC-seq modules, respectively. Next, we used Seurat’s anchor-based integration method to integrate snATAC and snRNA modalities based on rlsi and rpca reductions, respectively, before using both modalities to construct a weighted nearest neighbors (WNN) graph projection of cells into a 2D UMAP representation. Samples that were pooled into the same 10X run were demultiplexed and subsequently treated as separate samples during data integration. Clustering using the Leiden algorithm as implemented in leidenalg^90^ was conducted on the WNN graph, and cell type identities were assigned to clusters based on expression of marker genes.

### Demultiplexing pooled multiome samples

To demultiplex pooled samples, we isolated genomic DNA from non-islet tissue using the PureLink Genomic DNA mini kit and genotyped using the Illumina Infinium Omni 2.5-8 assay by the UCSD Institute for Genomic Medicine. During dithizone staining and islet picking, non-islet cells were collected separately from stained islets, washed with 1X HBSS, pelleted at 500rcf for 5 min, and snap frozen with liquid nitrogen until genomic DNA extraction. Genotypes were called using GenomeStudio (v2.0.4) with default settings and used PLINK^91^ to filter out rare variants with MAF <0.01 in the Haplotype Reference Consortium panel r1.1 and ambiguous alleles with MAF >0.4. Genotypes were then imputed into the HRC r1.1 panel using the Michigan Imputation Server with minimac4. Genotypes with high imputation quality (R2>0.3) were used to demultiplex pooled snMultiome samples using Demuxlet^92^ with default settings. Any cells not confidently assigned to a donor were removed from analysis.

### Pseudobulk differential expression analysis

We excluded all multiome samples from the Wang et al. study^24^ for this analysis. We pseudobulked gene expression by sample and cell type and used the design ∼donor + treatment for analysis. We used the Wald test with a local fit to determine differential gene expression. We considered cCREs significant at padj<0.05.

### Pseudobulk differential chromatin accessibility analysis

We excluded all multiome samples from the Wang et al. study^24^ for this analysis. We pseudobulked counts of fragments mapping to the union set of CREs in each cell type by sample by multiplying average expression by number of cells per sample. Pseudobulk count matrices for each cell type were input to DESeq2 with the design ∼donor + treatment. We used the Wald test with a local fit to calculate differential accessibility between cCREs. We considered genes significant at padj<0.05.

### Enrichment tests comparing bulk and pseudobulk differential expression

We defined the ‘universe’ of genes as the set of 15,301 unique gene symbols with adjusted p-values in differential analysis from both pseudobulk beta cells and bulk islets. For each treatment condition and for alpha and beta cells separately, we used the Fisher’s exact test to calculate pairwise enrichment for sets of significantly up- and down-regulated genes and used the BH test correction to correct for the total number of pairwise comparisons performed.

### Enrichment tests comparing bulk and pseudobulk differential accessibility

We annotated using bedtools overlap a set of 111,404 pairs of regions in which a cCRE from bulk islets overlapped with a cCRE from snATAC. Then, for each pair of regions and treatment conditions, and for alpha and beta pseudobulk results separately, we annotated whether the multiome cCRE was differentially up- or down-regulated in our pseudobulk results and whether the bulk islet cCRE was differentially up- or down-regulated. We used the Fisher’s exact test to determine pairwise enrichment of gene sets and performed the BH test correction for the total number of pairwise comparisons using the R base function ‘p.adjust’.

### MAGMA GWAS enrichment

We downloaded summary GWAS statistics for T2D^93^ and fasting glycemia^78^. We assigned SNPs to genes based on their genomic coordinates using the NCBI GRCh38/hg38 SNP positions. Gene-based association statistics were computed using a multiple linear principal components regression model and using the European-ancestry reference panel from the 1000 Genomes Project^94^ to account for linkage disequilibrium. We performed one-sided competitive gene-set analysis using MAGMA^33^ to test custom gene sets for enrichment of trait-related genes and used the Benjamini-Hochberg correction to adjust P-values to account for multiple tests.

### Ranked pathway enrichment analysis

We used fgsea to conduct ranked enrichment analysis of the GO version 2023.1 database on DESeq2^84^ results using the Wald test statistic to rank genes and included all genes with a non-NA p-value for this analysis. The GO v.2023 database^95^ used for analysis was obtained from MSigDB.

### ChromVAR motif analysis

We did not include ND control samples from the Wang et al study^24^ in this analysis. For this analysis, we subsampled snATAC and multiome-snATAC to a maximum of 3,000 cells per cell type for alpha and beta cells, with separate objects for each cell type, and added GC bias using the ‘BSgenome.Hsapiens.UCSC.hg38’ library from Bioconductor for genome sequence input. We used the motifmatchr package to annotate union peaks for motif occurrences using the JASPAR database and ran chromVAR^96^ to compute motif activity in individual cells. We used Seurat’s FindMarkers function with rowMeans to average motif activity in each cell type to compare motif activity scores in cyt- or TGN-treated cells compared to untreated cells. The average difference between treated and control cells was plotted for each transcription factor.

### Defining enhancer-gene links in beta cells

We concatenated ATAC fragments from all multiome and snATAC-seq beta cells into a single pseudobulk beta cell bam file used as activity input for ABC^28^. We used the union set of peaks from snATAC as potential enhancer regions. We used the power law to predict contact frequency and for enhancer scoring in our final analyses. ABC’s predicted ‘enhancer’ regions were mapped back to union peaks with bedtools overlap. In addition to cCRE-gene links predicted from ABC, we used bedtools overlap to assign target genes to cCREs based on overlap with gene body coordinates from NCBI38/GRCh38 extended 10kb the upstream and downstream direction.

### Beta cell WGCNA

We used the package hdWGCNA^97^ for this analysis with default settings unless otherwise indicated. In brief, we subset the islet snRNA-seq object to only beta cells for this analysis, then constructed and scaled expression of metacells consisting of up to 25 cells originating from the same sample. We then subset to only beta cells from the Wang et al^24^ study and constructed a signed network from the 15,000 most variable genes with using a soft power threshold of 6.

### Differential network expression analysis

We used the ‘FindDMEs’ method from hdWGCNA^97^ which leverages the Wilcoxon test to assess network expression in cytokine- or thapsigargin-treated beta against untreated controls generated from the same study. We also used the FindDMEs framework to test for differential network expression in beta cells from T2D donors compared to cells from ND donors. For this analysis, we did not include untreated samples generated in our study in the comparison of T2D vs. control, and we did not include ND donors from the Wang et al. study^24^ in comparisons of thapsigargin and cytokines vs. control. A brief description of the samples used for all DME analysis is included in **Table 18**.

### Pathway enrichment in hdWGCNA networks

Sets of genes assigned to each network were tested for pathway enrichment using GOSeq with the background set defined as 13,918 unique gene symbols with non-NA adjusted p-values in both pseudobulk beta cell gene expression cyt vs control and TGN vs control differential gene expression analyses, which included all genes assigned a network.

### Motif enrichment in hdWGCNA network enhancers

cCREs linked to candidate target genes, as described above, were grouped corresponding to target gene membership in co-expression networks. Notably, while gene networks are mutually exclusive, some cCREs target multiple genes, potentially in different networks. These sets of cCREs were used for motif analysis using HOMER as described above, using the set of all cCREs in the snATAC/multiome peak set as the background.

